# Ampyrone (4-Aminoantipyrine) is a Direct Agonist of Human Tyrosinase and Potential Therapeutic for Oculocutaneous Albinism and Disorders of Hypopigmentation

**DOI:** 10.1101/2025.10.13.682036

**Authors:** Monika B. Dolinska, Yuhong Wang, Nathan P. Coussens, Vijay K. Kalaskar, Zuhal Eraslan, Samuel J. Grondin, Joseph Bonica, Sarah Toay, Matthew D. Hall, Min Shen, Matthew Boxer, Qiuying Chen, Steven S Gross, Nabeel Attarwala, Yingyos Jittayasothorn, Ramakrishna P. Alur, Dhyanam Shukla, Robin Kee, Charles DeYoung, Cuilee Sha, David R. Adams, Stacie Loftus, Tiziana Cogliati, Yuri V. Sergeev, Jonathan H. Zippin, Brian P. Brooks

## Abstract

Significant loss of pigmentation can increase visual disability, skin cancer risk, and psychosocial stress. Tyrosinase (TYR) catalyzes the first and rate-limiting step of melanin synthesis. Inhibitors of TYR are well established and are currently used in clinical settings; however, there is a dearth of direct activators of TYR. Here, using a unique human TYR construct, high-throughput screening, and computational analysis techniques, we identified ampyrone as a TYR activator. Ampyrone increased the *in vitro* catalytic activity of the intramelanosomal domain of human TYR (hTYR) and its hypomorphic variant, P406L, a cause of oculocutaneous albinism type 1B (OCA1B). Moreover, ampyrone induced melanin synthesis in both wild-type and OCA1B human melanocytes, as well as 3-dimension (3D) human skin cultures. Our results reveal ampyrone as a lead compound for first-in-class TYR activators, potentially accelerating the discovery of novel therapies for patients with genetic and acquired diseases of hypopigmentation.

## Introduction

Pigmentation of the hair, eyes, and skin is not only a notable phenotypic characteristic, but it also protects from ultraviolet (UV) radiation and affects visual acuity. Disorders of pigmentation can be acquired (e.g., post-inflammatory, metabolic abnormalities) or genetic (e.g., oculocutaneous albinism (OCA)) (*1-3*). In many acquired instances, these pigmentary disorders are transient in nature, but, in some individuals, the effects can last for many years or never resolve. Cutaneous hypopigmentation increases skin cancer risk and can lead to negative psychosocial effects, including social stigmatization, reduced self-esteem, and heightened self-consciousness (*4, 5*). Individuals with OCA may present with an uncorrectable visual disability, including legal blindness. Despite the morbidity associated with hypopigmentation of the skin and eyes, there are limited therapeutic approaches.

Pigmentation in animals is primarily a reflection of melanin concentration. Melanin is a polymer formed in a specialized organelle called the melanosome, found exclusively in two cell types: (1) the neural crest derived melanocytes present in the hair, epidermis, meninges of the brain, stria vascularis of the ear, and the choroid and iris of the eye and (2) the neuroectoderm-derived pigmented epithelial cells, such as retinal pigment epithelium (RPE) cells of the eye (*6*). Tyrosinase (TYR) catalyzes the initial and rate-limiting steps of melanin production. Over the last few decades, numerous cosmeceutical and pharmaceutical companies have used non-vertebrate TYR to screen for small molecules that alter melanin synthesis, leading to the discovery of various TYR inhibitors (*7, 8*); however, activators of melanin synthesis have remained elusive (*9*). Direct activators of TYR would have significant clinical utility in almost all forms of albinism and could represent a new therapeutic approach for improving pigmentation in those with fair skin and increased risk of skin cancer (*10-12*).

We recently described the expression and purification of the human recombinant intramelanosomal domain of TYR (hTYR, hTYR^WT^) and OCA1B-related hypomorphic variants (hTYR^P406L^) (*13-15*). Here, we developed a high-throughput screening (HTS) approach using the hTYR^WT^ protein to simultaneously identify its activators and inhibitors. After screening over 34,000 compounds, we identified seven potential activators and 65 potential inhibitors (32 of which were novel). Four activators and six novel inhibitors were further examined with detailed enzymology. Among the potential activators, ampyrone (4-aminoantipyrine; IUPAC, 4-Amino-2,3-dimethyl-1-phenyl-3-pyrazol-5-one) increased the catalytic activity of both hTYR^WT^ and hTYR^P406L^. Computational analysis indicates that ampyrone enhances catalytic activity and may help restore hTYR^P406L^ by improving structural stability and reorganizing active site geometry, including copper-histidine coordination and substrate placement, resulting in a more rigid and energetically stable active site. Using a newly developed liquid chromatography-mass spectroscopy (LC-MS) based tyrosine tracing technique we observed increased melanin synthesis in live human normal and OCA1B melanocytes within minutes of ampyrone treatment. Finally, ampyrone also increased epidermal pigmentation in a three-dimensional (3D) skin model. Overall, our findings suggest that ampyrone could be a promising therapeutic lead compound for developing a new class of TYR agonists to treat albinism and other hypopigmentation disorders and to protect fair-skinned individuals from UV damage.

### High-throughput screening (HTS) identified potential activators and inhibitors of hTYR^WT^

hTYR^WT^ and hTYR^P406L^ enzymes were purified and optimized for primary and/or secondary screening of small molecule activators or inhibitors (**Supplemental Figure S1, S2**). Previous *in vitro* experiments confirmed that the hTYR^P406L^ variant retains 35% of hTYR^WT^ specific activity (*14*).

hTYR catalyzes the oxidation of mono- and diphenols (L-tyrosine and L-DOPA, respectively) to their corresponding quinones, which, in turn, produce dopachrome, an orange/brown product that absorbs light at 475 nm and can be measured spectrophotometrically. We developed a miniaturized 1536-well kinetic assay for hTYR^WT^ diphenol oxidase activity that was optimized to identify activators and inhibitors from chemical libraries that were unaffected by vehicle and exhibited Z’-factor values > 0.8 (**Supplemental Figure S3A-C**). A primary HTS was conducted on 34,051 compounds with a 4% average coefficient of variation for the vehicle-treated baseline control and an average Z’-factor of 0.74 (activator) and 0.79 (inhibitor) (**Supplemental Figure S3D-F, 4A**). Among the hits from the primary screen, 115 compounds with inhibitory activity were identified, of which six were selected, based on their preliminary activity, for evaluation in a confirmatory *in vitro* assay (**Supplemental Figure S4B, C, and Supplemental Table S1**). The diphenol oxidase activity of hTYR^WT^ was inhibited in a concentration-dependent manner by all six compounds, 4-anilinophenol, idronoxil, anethole trithione, pestanal (fenaminosulf), 3’,4’-dihydroxyflavone, and 6-thioguanine, with IC_50_ values of 3.20, 29.13, 1.45, 8.12, 9.22, and 540 µM, respectively (**Supplemental Figure S5A)**. A total of nine compounds were identified as potential activators, of which four were confirmed using the same secondary *in vitro* assay applied for inhibitors (**Supplemental Figure S4B, C, and Supplemental Table S1**). Of the other five potential activators, phloretin and brazilin were excluded due to prior characterization as TYR inhibitors (*16, 17*), 4,4’-Thiodianiline because of its known carcinogenicity, and the two compounds from the Genesis library were unavailable. Of the four confirmed hits, hydroxytacrine maleate exhibited a modest, concentration-dependent increase in absorbance at 475 nm over time, while sennoside B showed a modest and non-concentration-dependent effect. In contrast, 4-dimethylaminoantipyrine had no effect at any of the tested concentrations (**Supplemental Figure S5B)**. Finally, a significant increase in hTYR^WT^ diphenol oxidase activity was observed in the presence of ampyrone (**Supplemental Figure S5C**).

### Ampyrone increases the enzymatic activity of hTYR^WT^ and hTYR^P406L^

Ampyrone demonstrated a concentration-dependent increase in the diphenol oxidase activity of both hTYR^WT^ and hTYR^P406L^ enzymes with EC_50_ values of 0.67 and 3.57 mM, respectively (**Supplemental Figure S5C, D)**. Incubation with 10 mM ampyrone increased the maximum reaction velocity (V_max_), Michaelis constant (K_m_), turnover number (k_cat_), and catalytic efficiency (k_cat_/K_m_) for both enzymes (**Figure 1A, Supplemental Figure S6A, and Table 1**). For hTYR^WT^, the k_cat_ increased by 73%, but the 91% increase in K_m_ resulted in a 13% decrease in catalytic efficiency. In the case of the hTYR^P406L^ variant, ampyrone increased the k_cat_ by 238% and K_m_ by 153%, resulting in a 35% increase in diphenol oxidase catalytic efficiency. Notably, ampyrone enhanced catalytic activity of the hTYR^P406L^ enzyme more substantially when expressed as a percentage of control. Although the affinity of L-DOPA, reflected by K_m_, decreased in both hTYR^WT^ and hTYR^P406L^ enzymes, this was offset by an overall increase in enzyme efficiency, leading to an increase in diphenol oxidase activity.

**Table 1.** Kinetic parameters of hTYR^WT^ and hTYR^P406L^ enzymes monophenolase and diphenol oxidase activity. Incubation with ampyrone increased the V_max_, K_m_, k_cat_, and k_cat_/K_m_. Data represent the mean ± SD from three replicate experiments.

| Kinetic Parameters | Monophenolase activity |  |  |  | Diphenol oxidase activity |  |  |  |
| --- | --- | --- | --- | --- | --- | --- | --- | --- |
|  | hTYR<sup>WT</sup> |  | hTYR<sup>P406L</sup> |  | hTYR<sup>WT</sup> |  | hTYR<sup>P406L</sup> |  |
|  | control | ampyrone | control | ampyrone | control | ampyrone | control | ampyrone |
| $V_{\max}$<br>(mM sec<sup>-1</sup>) | 0.19 ± 0.06 | 0.30 ± 0.04 | 0.15 ± 0.01 | 0.46 ± 0.02 | 0.21 ± 0.02 | 0.36 ± 0.01 | 0.10 ± 0.01 | 0.32 ± 0.01 |
| $K_m$<br>(mM) | 0.11 ± 0.06 | 0.05 ± 0.01 | 0.12 ± 0.02 | 0.04 ± 0.01 | 0.22 ± 0.06 | 0.42 ± 0.02 | 0.40 ± 0.01 | 1.01 ± 0.08 |
| $k_{\text{cat}}$<br>(sec<sup>-1</sup>) | 105.14 ± 35.12 | 168.35 ± 21.72 | 83.71 ± 6.32 | 258.52 ± 10.21 | 118.35 ± 10.17 | 204.06 ± 4.50 | 54.51 ± 1.43 | 184.34 ± 3.92 |
| $K_{\text{cat}}/K_m$<br>(mM sec<sup>-1</sup>) | 990.50 ± 182.43 | 3198.10 ± 54.98 | 707.96 ± 66.94 | 6285.45 ± 217.72 | 555.41 ± 134.38 | 484.39 ± 29.45 | 135.56 ± 0.29 | 182.28 ± 10.17 |

**Figure 1.**
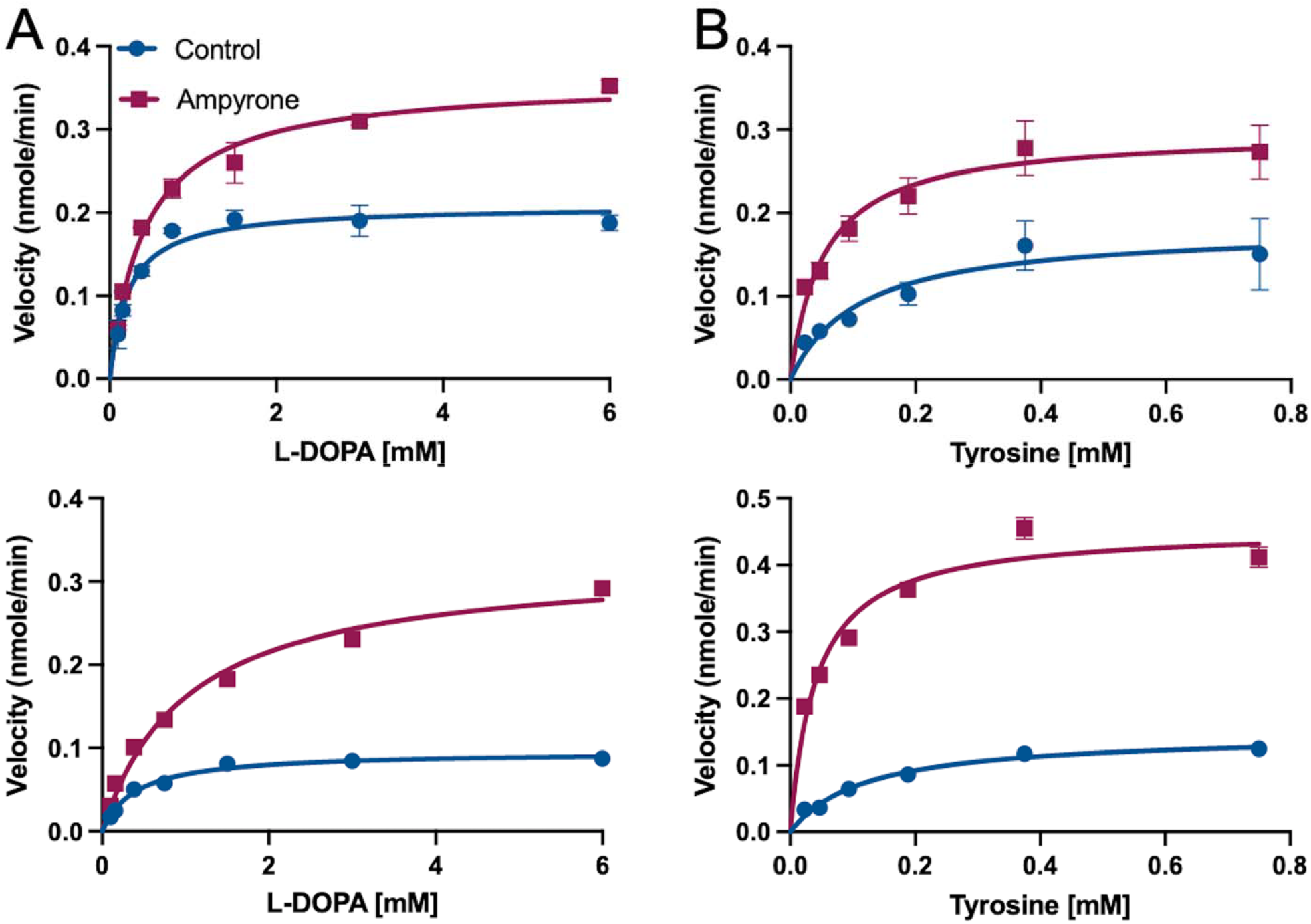
Effect of 10 mM ampyrone on the diphenol oxidase and monophenolase activity of hTYR^WT^ and hTYR^P406L^. Michaelis–Menten plots of diphenol oxidase (A) and monophenolase (B) activity of hTYR^WT^ (*top)* and hTYR^P406L^ (*bottom)* enzymes measured at 37°C in the presence (*red)* or absence (control, *blue)* of 10 mM ampyrone using increasing concentrations of L□DOPA or tyrosine, respectively. Curves represent nonlinear fits to the Michaelis–Menten equation generated in GraphPad Prism 10. Data represent the mean +/-SD from three replicate experiments.

Further kinetic analyses showed that ampyrone also significantly increased monophenolase activity in both hTYR^WT^ and hTYR^P406L^ enzymes (**Figure 1B, Supplemental Figure S6B, and Table 1**). In the hTYR^WT^, ampyrone increased V_max_ by 58%, decreased K_m_ by 55%, and enhanced k_cat_ by 60%, resulting in a 222% increase in catalytic efficiency. The response was even more pronounced in hTYR^P406L^: ampyrone increased V_max_ by 207%, decreased K_m_ by 67%, and elevated k_cat_ by 209%, resulting in a 788% increase in catalytic efficiency. These findings indicate that ampyrone strongly boosts monophenolase efficiency, especially in hTYR^P406L^, while also enhancing diphenol oxidase activity.

### Modeling of ampyrone effects on hTYR

One hindrance to understanding hTYR enzymology is the lack of a TYR x-ray crystal structure. Therefore, to examine how ampyrone binding leads to an increase in enzyme activity, we performed molecular modeling of hTYR based on the crystal structure of the closely related enzyme hTYRP1 (*18*). Ampyrone was docked to models of both the hTYR^WT^ and hTYR^P406L^. Several binding positions outside the active site were shared between the two proteins. The top three binding sites were selected for further molecular dynamic (MD) simulations (**Supplemental Figure S7**). In both enzyme variants, one ampyrone molecule consistently bound near residues P205-T226. A second molecule bound at the interface between the signal peptide, the 150-170 loop, and the β-sheet 200 region. The third binding site differed between the two variants: in the hTYR^WT^ model, ampyrone bound to the loop connecting β-sheets within the Cys-rich subdomain, whereas in the hTYR^P406L^ model, it occupied a cavity separating the Cys-rich and catalytic subdomains.

We next sought to understand how ampyrone binding affects the conformation and flexibility of hTYR residues using MD simulations. The root-mean square deviation (RMSD) measures the average change in atomic positions between two structures over time, providing an overall view of structural stability. In contrast, the root mean-square fluctuation (RMSF) reflects the average positional fluctuation of individual residues, offering insight into local flexibility. Ampyrone binding had minimal impact on the RMSD profile of hTYR^WT^ but notably reduced the elevated RMSD observed in hTYR^P406L^, bringing it closer to the wild-type profile (**Supplemental Figures S8-S9**). The RMSF plot revealed reduced movement upon ampyrone binding to both hTYR^WT^ and hTYR^P406L^, with the latter displaying a more pronounced effect. However, ampyrone did not restore coordinated motions within helices I172-S184 and P205-T226 as reflected by the lack of recovery in dynamical cross-correlation matrix (DCCM) patterns, which quantify how pairs of residues move in a correlated or anti-correlated manner throughout MD simulations (**Supplemental Figure S10**).

Additionally, the presence of ampyrone resulted in the rearrangement of active site residues in hTYR, including the six copper-coordinating histidine residues, impacting the binding position of L-tyrosine (**Figure 2A, B**). Distances and coordination angles between copper atoms and H202, H211, and H363 shifted substantially and consistently for both hTYR^WT^ and hTYR^P406L^ (**Supplemental Figures S11-S12**). These structural perturbations in the presence of ampyrone were further demonstrated by porcupine plots, which illustrate both the direction and extent of residue movements along the dominant motion pathway during the simulation. These plots showed increased mobility of residues near copper sites (e.g., F207, H202, E203), while more distal residues remained relatively stable (**Figure 2C-D**). The direction of movement, however, appears to be residue specific. To assess how ampyrone affects active site conformational dynamics, we performed free energy landscape (FEL) analysis using distances between Cß atoms of V377 and the side chains of two catalytically important residues: F347 (CZ) and K334 (NZ), as reaction coordinates. Ampyrone binding reduced conformational heterogeneity in both hTYR^WT^ and hTYR^P406L^, as shown by narrower and deeper energy minima, indicating a more rigid and energetically stabilized active site (**Figure 2E-H**). Modeling predictions that ampyrone improves hTYR protein stability were experimentally confirmed using a urea denaturation approach (**Supplemental Figure S13**).

**Figure 2.**
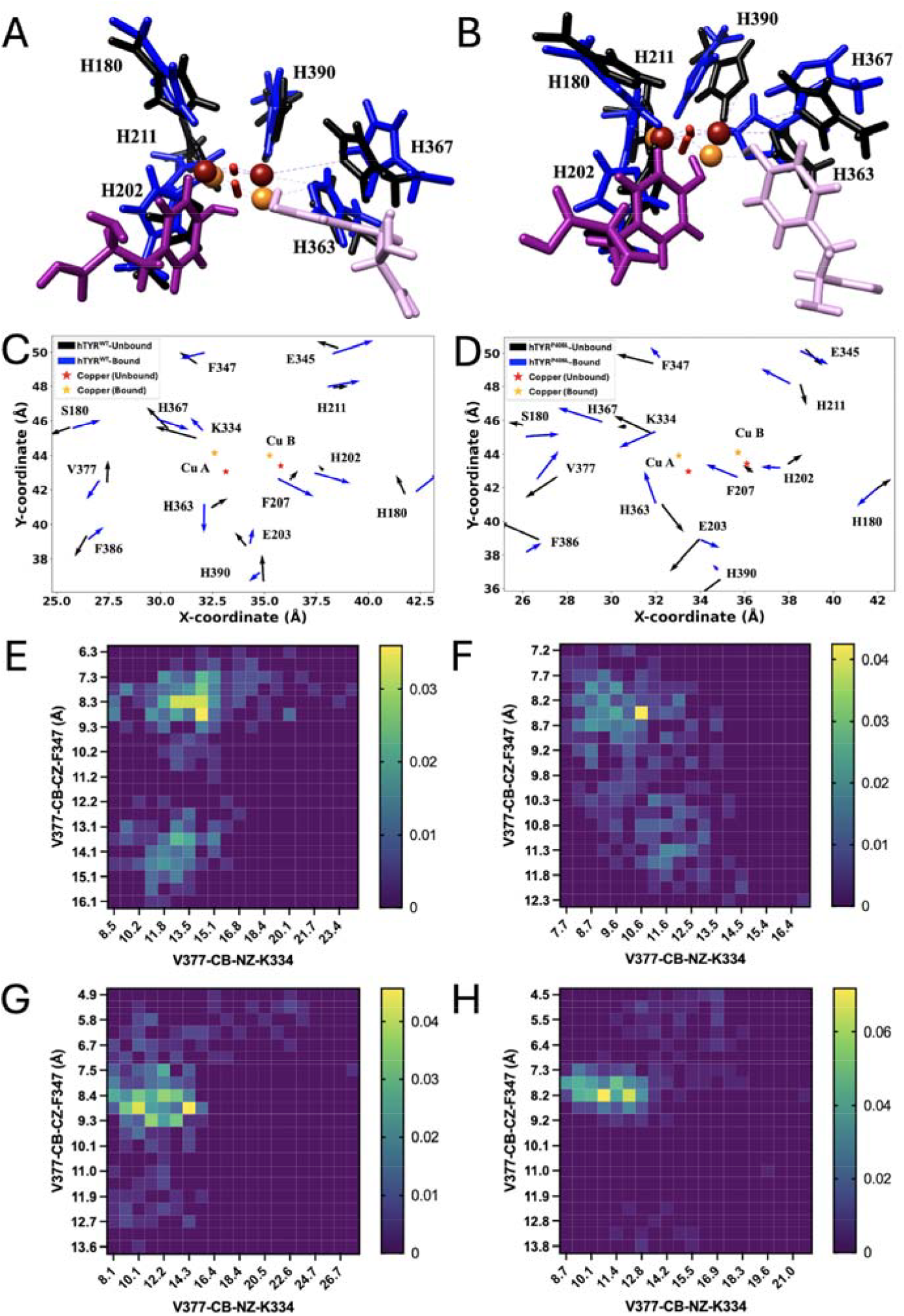
Computational modeling of ampyrone effects in hTYR^WT^ and hTYR^P406L^. The active site in the absence (Protein/Black, Substrate/Light purple, Cu/Red, O_2_/Red) and presence (Protein/Blue, Substrate/Dark purple, Cu/Orange, O_2_/Red) of ampyrone shows movements in the six catalytic histidine residues for both hTYR^WT^ (A) and hTYR^P406L^ (B) when binding L-tyrosine. Coppers A and B (CuA and CuB) can be read left to right in all panels. Porcupine plots display the movements of the alpha carbon atoms of select active site residues in the absence and presence of ampyrone for hTYR^WT^ (C) and hTYR^P406L^ (D). Free energy landscapes mapping the distance V377-CB-CZ-F347 and V377-CB-NZ-K334 in angstroms (Å) for hTYR^WT^-unbound (E), hTYR^WT^-bound (F), hTYR^P406L^-unbound (G), hTYR^P406L^-bound (H).

### Ampyrone increases eumelanin production in human normal and OCA1B melanocytes

Traditional methods of measuring melanin rely on the accumulation of polymerized melanin and the analysis of total cellular melanin by quantifying degradation products by HPLC(*19*). However, this approach cannot capture immediate changes in melanin synthesis. We recently developed an LC-MS method for measuring melanin synthesis that utilizes ^13^C-labeled tyrosine, thereby allowing for the detection of melanin synthetic intermediates over time (*20*). Using this method, we found that while ampyrone did not affect tyrosine uptake into melanocytes, it induced an increase in the two main eumelanin synthetic intermediates, 5,6-dihydroxyindole-2-carboxylic acid (DHICA) and 5,6-dihydroxyindole (DHI), within 20 minutes, the earliest time point we can measure (**Figure 3A**). Ampyrone treatment induced a further increase in DHICA and DHI at one hour, which then plateaued at three hours, suggesting an equilibrium between new synthesis and polymerization into melanin (**Figure 3A**). We next examined the effect of ampyrone on melanin synthesis over multiple days in wild-type and OCA1B human melanocytes by measuring changes in side-scatter using flow cytometry and found that ampyrone treatment for 48 hours increased melanin synthesis (**Figure 3B and Supplemental Figure S14A, B**). These findings indicate that ampyrone treatment may be an effective approach for increasing melanin synthesis in melanocytes with both wild-type and defective TYR activity.

**Figure 3.**
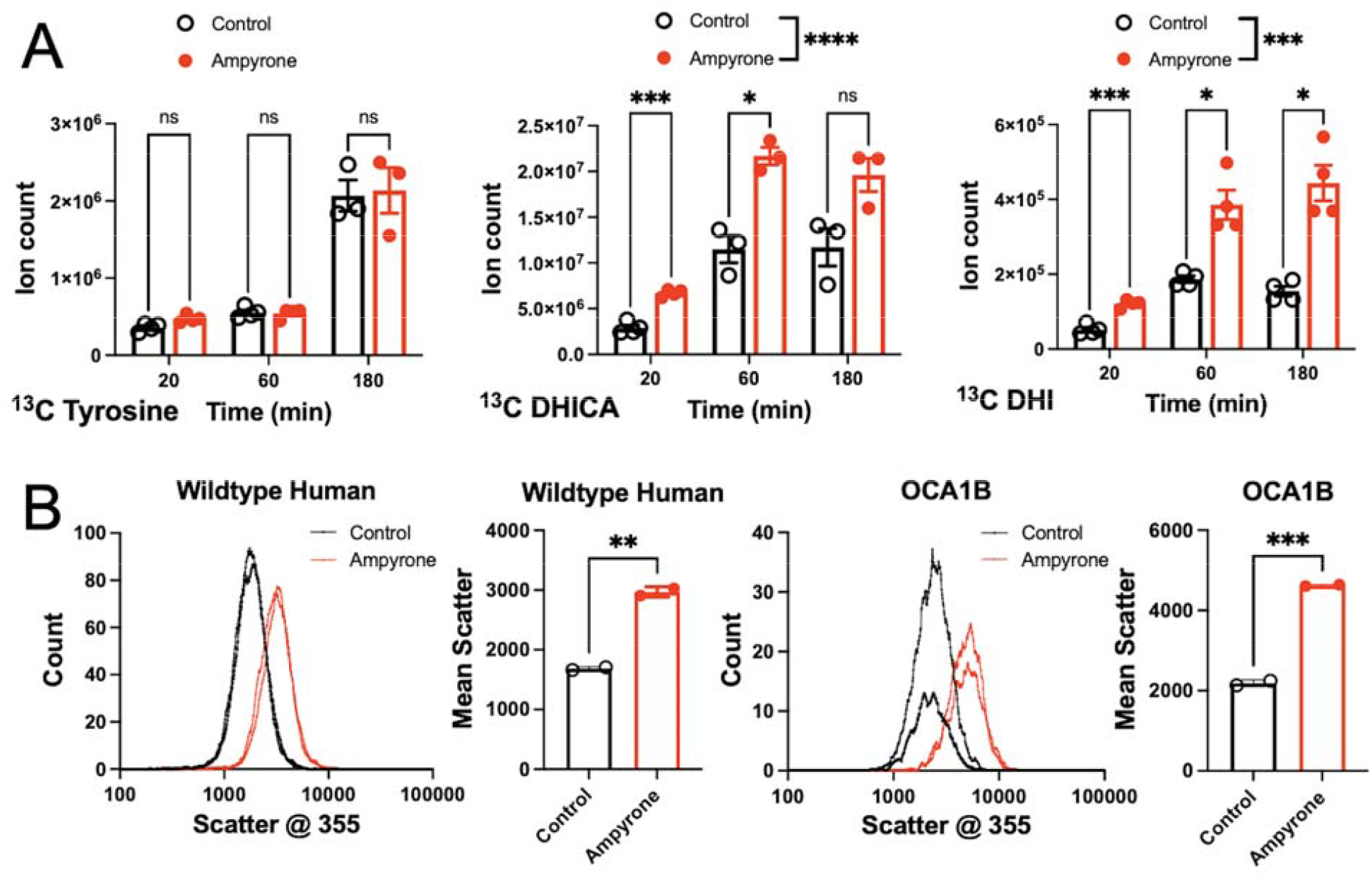
Ampyrone induces melanin synthesis in melanocytes. A) LC-MS tracing of ^13^C-tyrosine metabolism in the absence (black) or presence of ampyrone (2mM, red). Ion counts detected over time of tyrosine (left), DHICA (middle), and DHI (right). ANOVA with post-hoc Tukey. N≥3. B) Flow cytometry of human wild-type (C4) and OCA1B-1125 melanocytes after three days in vehicle (control, black) or ampyrone (2mM, red). The left panels show histograms of cell counts at different amounts of scatter at 355 nm. The right panels show the mean scatter at 355 nm. Raw scatter plots shown in Figure S14. Student’s t-test, unpaired (N≥2), ns = not significant, *, P<0.05. **, P<0.01. ***, P<0.001.

### Ampyrone increases pigmentation in a human 3D epidermal model

Skin pigmentation is primarily driven by complex interactions between melanocytes and keratinocytes, the principal cellular components of the epidermis. We next assessed whether ampyrone treatment increases epidermal pigmentation in a human 3D epidermal culture system consisting of co-cultured normal, human-derived epidermal keratinocytes (NHEK) and melanocytes (NHM). This artificial human skin model is widely used to study epidermal pigmentation *in vitro* (*21, 22*). Ampyrone treatment (200µM) increased epidermal and melanocyte pigmentation over 21 days of treatment (**Figure 4A, B**). Histological staining followed by quantitation of epidermal pigmentation further confirmed that ampyrone treatment increased epidermal pigmentation without any evidence of epidermal toxicity (**Figure 4C, Supplemental Figure S15A, B**), suggesting that ampyrone treatment could be used to induce pigmentation in human epidermis.

**Figure 4.**
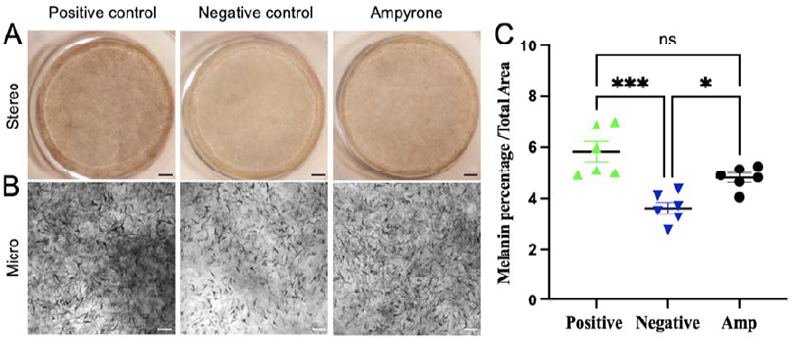
Ampyrone treatment increases pigmentation of 3D human skin culture. A) Stereo microscopic (Stereo (10X) images taken from representative positive and negative control treated and ampyrone-treated (200µM) MelanoDerm™ samples. B) Inverted microscopic (Micro, 10X) images showing representative images of melanocyte pigmentation following treatment with ampyrone (200µM), positive and negative control media. C) Quantitation of Fontana-Masson staining (melanin) expressed as a percentage across the entire section area. N=2, three replicates. ANOVA post-hoc Tukey. *, P<0.05. ***, P<0.001. ns=not significant. Scale bar in panel A: 1000 µm and B: 100 µm. These data are representative of an experiment repeated three times.

## Discussion

Millions of people every year can suffer from insufficient melanin synthesis in the skin and eyes, due to genetic or acquired mechanisms, which can decrease visual acuity, increase the risk of skin cancer, and accelerate signs of aging. In OCA we posit that restoring ocular melanin early enough in life, when foveal development is proceeding, could result in improved visual function (*23*), and that improving skin pigmentation could reduce the risk of skin cancer, and alleviate psychosocial stress and associated morbidity. While some effective and safe drugs that inhibit melanin synthesis by targeting TYR are FDA-approved, there are currently no FDA-approved drugs that directly stimulate TYR.

To address this unmet medical need, we developed an HTS approach using multiple enzymatic activities of a uniquely generated hTYR to identify both inhibitors and activators. Using this novel approach, we identified numerous inhibitors of hTYR with structurally novel chemical backbones (**Supplemental Table S1**) and, importantly, several activators of both hTYR^WT^ and the hTYR^P406L^ disease variant with previously undescribed chemical structures (**Supplemental Table S1**). These data highlight the power of our screening approach, which could be applied to other hTYR disease-causing variants, and its potential to reveal novel regulators of pigmentation. When using traditional methods of measuring in-cell melanin synthesis, background melanin levels prevent the detection of immediate changes. In this manuscript, we also highlighted the power of LC-MS-based ^13^C-tyrosine tracing as a method for the discovery and validation of activators or inhibitors of hTYR. Tyrosine tracing can detect changes in melanin synthesis within 20 minutes of ampyrone treatment, allowing for cell-based screening of drugs with near cell lysate level speed. We predict this approach will be effective for identifying melanin synthesis-modifying drugs that target melanosome pH and other melanosome biology, which cannot be modeled in a protein lysate.

We focused our efforts on characterizing the effects of ampyrone on hTYR^WT^ and the common OCA1B-associated hTYR variant P406L. Compounds related to ampyrone are already approved for use in humans, which would potentially facilitate its translation into human trials. We found that ampyrone potently stimulates the catalytic efficiency of both hTYR^WT^ and hTYR^P406L^ enzymes, targeting both mono- and diphenol oxidase activities. Ampyrone treatment showed no overt toxicity in human melanocytes in culture and in human 3D skin model cultures, even after three weeks of exposure, suggesting that it could serve as a lead compound for the development of safe and effective TYR agonists. Our modeling predicts that ampyrone transiently binds to a diverse array of pockets on the surfaces of both the hTYR^WT^ and hTYR^P406L^ protein structures. Interestingly, these areas are relevant for coordinated movements within the bundle of core alpha helices (*24*), potentially suggesting that ampyrone primarily targets these flexible regions of the hTYR protein. This is further supported by the observed reduction in movement within the regions bordering the junction of the cysteine-rich and catalytic subdomains. As anticipated, the hTYR^P406L^ protein exhibits a more significant reduction in residue fluctuations over time induced by ampyrone. This provides computational evidence that ampyrone may help restore TYR function in mutants by enhancing structural stability, in addition to boosting hTYR^WT^ enzyme activity. Ampyrone also induces significant rearrangement of catalytic residues in both hTYR^WT^ and hTYR^P406L^. These changes affect the distances and coordination angles between the six copper-coordinating histidines with histidine-specific alterations and, in some instances, increased flexibility that may enhance L-tyrosine binding and catalytic efficiency. Additionally, ampyrone influences residues further from the coppers, promoting reduced conformational flexibility in positions such as K334, F347, and V377. These residues likely play integral roles in substrate binding rather than direct catalysis (*25*). Thus, ampyrone may induce both hTYR^WT^ and hTYR^P406L^ to adopt catalytically relevant active site conformations more frequently, thereby enhancing substrate binding.

Finally, our study demonstrates that ampyrone significantly enhances pigmentation in a human 3D epidermal model. Notably, this increase in pigmentation occurred without any signs of epidermal toxicity, indicating the compound’s safety. These data further support the use of ampyrone as a lead molecule for the development of therapeutics for patients with hypopigmentation of the skin or eyes.

In summary, using newly designed HTS and LC-MS approaches to screen for activators of our recently developed hTYR protein, we identified a new class of hTYR agonists with the therapeutic potential to treat patients with disorders of hypopigmentation, including OCA, which is currently an orphan disease. We predict that the development of therapeutics capable of increasing human pigmentation have the potential to reduce skin cancer risk, improve visual acuity, and alleviate the psychosocial morbidity of patients with deficient melanin synthesis.

## Acknowledgments

We thank members of the Zippin and Brooks lab for critical reading of the manuscript. We would like to thank the staff of the NEI Histology Core for their technical help in sectioning and staining specimens. **Funding:** J.H.Z was funded in part by NIAMS (1 R01 AR077664-01A1) and the National Organization for Albinism and Hypopigmentation Established Researcher Program Grant. S.J.G was funded in part by NCI (T32 CA062948). This research was supported in part by the Intramural Research Program of the National Institutes of Health (NIH), National Eye Institute and the National Center for Advancing Translational Sciences. The contributions of the NIH authors are considered works of the United States Government. The findings and conclusions presented in this paper are those of the author(s) and do not necessarily reflect the views of the NIH or the U.S. Department of Health and Human Services.

## Author Contributions

M.B.D., Y.V.S., N.P.C., J.H.Z., and B.P.B. designed the experiments. M.B.D., S.T., Z.E., S.J.G., J.B., Y.W., and N.P.C. generated the Figures. Q.C., S.S.G., D.A., and S.L. generated critical reagents or assisted with LC-MS experiments. M.B.D., J.H.Z., and B.P.B. wrote the manuscript with all authors providing feedback.

## Materials and Data Sharing

All cell lines, drugs used, antibodies and other reagents used in the study are freely available from commercial sources or from publicly available biorepositories as detailed in the manuscript. Raw screening data, LC-MS data, and molecular modeling data will be made freely available to all interested parties. Human TYR cDNA plasmids will be provided upon request.

## References

1. M. G. Thomas, J. Zippin, B. P. Brooks, in GeneReviews((R)), M. P. Adam et al., Eds. (Seattle (WA), 1993).

2. L. Krueger, A. Saizan, J. A. Stein, N. Elbuluk, Dermoscopy of acquired pigmentary disorders: a comprehensive review. Int J Dermatol 61, 7–19 (2022).

3. M. D. Saleem, E. Oussedik, J. J. Schoch, A. C. Berger, M. Picardo, Acquired disorders with depigmentation: A systematic approach to vitiliginoid conditions. J Am Acad Dermatol 80, 1215–1231 e1216 (2019).

4. H. Fournier, N. Calcagni, F. Morice-Picard, B. Quintard, Psychosocial implications of rare genetic skin diseases affecting appearance on daily life experiences, emotional state, self-perception and quality of life in adults: a systematic review. Orphanet J Rare Dis 18, 39 (2023).

5. C. Garbe et al., Skin cancers are the most frequent cancers in fair-skinned populations, but we can prevent them. Eur J Cancer 204, 114074 (2024).

6. M. Snyman, R. E. Walsdorf, S. N. Wix, J. G. Gill, The metabolism of melanin synthesis-From melanocytes to melanoma. Pigment Cell Melanoma Res 37, 438–452 (2024).

7. H. Ando, H. Kondoh, M. Ichihashi, V. J. Hearing, Approaches to identify inhibitors of melanin biosynthesis via the quality control of tyrosinase. J Invest Dermatol 127, 751– 761 (2007).

8. M. A. Baber, C. M. Crist, N. L. Devolve, J. D. Patrone, Tyrosinase Inhibitors: A Perspective. Molecules 28, (2023).

9. S. Zolghadri et al., A comprehensive review on tyrosinase inhibitors. J Enzyme Inhib Med Chem 34, 279–309 (2019).

10. C. Niu, H. A. Aisa, Upregulation of Melanogenesis and Tyrosinase Activity: Potential Agents for Vitiligo. Molecules 22, (2017).

11. S. Guan, W. Su, N. Wang, P. Li, Y. Wang, A potent tyrosinase activator from Radix Polygoni multiflori and its melanogenesis stimulatory effect in B16 melanoma cells. Phytother Res 22, 660–663 (2008).

12. S. Guan, W. Su, N. Wang, P. Li, Y. Wang, Effects of radix polygoni multiflori components on tyrosinase activity and melanogenesis. J Enzyme Inhib Med Chem 23, 252–255 (2008).

13. M. B. Dolinska et al., Albinism-causing mutations in recombinant human tyrosinase alter intrinsic enzymatic activity. PLoS One 9, e84494 (2014).

14. M. B. Dolinska et al., Oculocutaneous albinism type 1: link between mutations, tyrosinase conformational stability, and enzymatic activity. Pigment Cell Melanoma Res 30, 41–52 (2017).

15. M. B. Dolinska, P. T. Wingfield, Y. V. Sergeev, Purification of Recombinant Human Tyrosinase from Insect Larvae Infected with the Baculovirus Vector. Curr Protoc Protein Sci 89, 6 15 11–16 15 12 (2017).

16. J. Chen et al., Phloretin as both a substrate and inhibitor of tyrosinase: Inhibitory activity and mechanism. Spectrochim Acta A Mol Biomol Spectrosc 226, 117642 (2020).

17. K. Mitani et al., Suppression of melanin synthesis by the phenolic constituents of sappanwood (Caesalpinia sappan). Planta Med 79, 37–44 (2013).

18. X. Lai, H. J. Wichers, M. Soler-Lopez, B. W. Dijkstra, Structure of Human Tyrosinase Related Protein 1 Reveals a Binuclear Zinc Active Site Important for Melanogenesis. Angew Chem Int Ed Engl 56, 9812–9815 (2017).

19. K. Wakamatsu, S. Ito, Advanced chemical methods in melanin determination. Pigment Cell Res 15, 174–183 (2002).

20. Q. Y. Chen et al., Measurement of Melanin Metabolism in Live Cells by [U-C]-L-Tyrosine Fate Tracing Using Liquid Chromatography-Mass Spectrometry. Journal of Investigative Dermatology 141, 1810–+ (2021).

21. H. Y. Kim et al., 2,4,6-Triphenyl-1-hexene, an Anti-Melanogenic Compound from Marine-Derived Bacillus sp. APmarine135. Mar Drugs 22, (2024).

22. M. Kim, C. S. Lee, K. M. Lim, Rhododenol Activates Melanocytes and Induces Morphological Alteration at Sub-Cytotoxic Levels. Int J Mol Sci 20, (2019).

23. I. F. Onojafe et al., Nitisinone improves eye and skin pigmentation defects in a mouse model of oculocutaneous albinism. J Clin Invest 121, 3914–3923 (2011).

24. S. Toay, Y. V. Sergeev, Genetic mutations disrupt the coordinated mode of tyrosinase’s intra-melanosomal domain. Protein Science 34, (2025).

25. M. B. Dolinska, Y. V. Sergeev, Molecular Modeling of the Multiple-Substrate Activity of the Human Recombinant Intra-Melanosomal Domain of Tyrosinase and Its OCA1B-Related Mutant Variant P406L. Int J Mol Sci 25, (2024).

