## Supplemental materials for "Ampyrone (4-Aminoantipyrine) is a Direct Agonist of Human Tyrosinase and Potential Therapeutic for Oculocutaneous Albinism and Disorders of Hypopigmentation"

**Materials and Methods**

***Expression and purification of WT and P406L intramelanosomal domains***

The recombinant intramelanosomal domain of WT hTYR (hTYR^WT^, residues 19 – 469) and the OCA1B-related P406L variant (hTYR^P406L^) were engineered with a His-tag, expressed using baculovirus, and produced in whole insect *Trichoplusia ni* larvae at Allotropic Tech, LLC (<https://allotropictech.com/>). The proteins were purified using immobilized metal affinity chromatography (IMAC) with a His-Trap crude 5 mL column, followed by gel-filtration (GF) chromatography on HiPrep 26/60 Sephacryl S-300 and Superose 12 10/300 GL columns (Cytiva), as previously described [1-3]. Fractions containing the protein of interest were concentrated using Amicon Ultra‐15/10,000 NMWL centrifugal filter units (Millipore Sigma). Protein concentrations were determined by measuring *A*_260/280 nm_ with a NanoDrop 2000c UV–Vis spectrophotometer (Thermo Fisher Scientific). The identities of hTYR^WT^ and hTYR^P406L^ proteins were confirmed by Western blot analysis using the Anti-TYR (T311) antibody (Santa Cruz Biotechnology, sc-20035).

***Miniaturized hTYR^WT^ diphenol oxidase activity assay and HTS***

Using HTS, ~34,000 compounds from the Genesis Diverse Chemical Library (Genesis), the National Center for Advancing Translational Sciences (NCATS), Pharmacologically Active Chemical Toolbox (NPACT), the NCATS Pharmaceutical Collection (NPC), and the Natural Products Library (NPL) were screened with purified hTYR^WT^. The diphenol oxidase activity assay was performed in black, medium-binding, clear flat-bottom 1536-well microplates (Greiner) with a final reaction volume of 8 µL. All steps were performed at room temperature. First, 6 µL of reaction buffer (50 mM sodium phosphate, 0.01% Triton X-100, pH 7.5) were dispensed into the 32 wells of column 1 (no enzyme, low activity control), 6 µL of reaction buffer containing 6 µM hTYR^WT^ [4.5 µM final] were dispensed into the 32 wells of column 2 (3-fold enzyme concentration, high activity control), and 6 µL of reaction buffer containing 2 µM hTYR^WT^ [1.5 µM final] were dispensed into the wells of columns 3 – 48. These solutions were dispensed with a BioRaptr workstation (Beckman Coulter). Next, 92 nL of dimethylsulfoxide (DMSO, vehicle control, columns 1 – 4) or library compounds (columns 5 – 48) were pin-transferred (Kalypsis) using a dip factor of 4. For the primary screen, library compounds from 10 mM stock solutions were tested at a final concentration of 115 µM. Following the transfer of vehicle or test compounds, the microplates were spun at 377 × *g* for 1 min and allowed to incubate for 30 min. Before the addition of substrate, absorbance was measured at 475 nm with a ViewLux uHTS microplate imager (PerkinElmer) to quantify any compound-related absorbance that was independent of hTYR^WT^ diphenol oxidase activity. The exposure time was 1 s with an excitation energy of 4,000, a readout speed of 10 µs, a 50X readout gain, and 2X image binning. The absorbance protocol included a 480 nm ± 10 nm (480/20) narrow band interference excitation filter and a clear infrared damping 400 nm – 750 nm emission filter. The hTYR^WT^ diphenol oxidase reaction was initiated by the addition of 2 µL of reaction buffer containing 1.72 mM L-DOPA (Sigma-Aldrich) [430 µM final] to all wells in columns 1 – 48. Immediately following substrate addition, the microplates were spun at 377 × *g* for 1 min and absorbance was measured at 30 s intervals for 20 min. Activity was measured as delta by comparing the absorbance at a final time point with that of the initial time point.

***Tyrosinase activation assays***

The diphenol oxidase activities of hTYR^WT^ and hTYR^P406L^ enzymes were measured spectrophotometrically using a SpectraMax i3 multi-mode detection platform, with data analyzed by SoftMax Pro software (version 6.5, Molecular Devices). Enzymes were incubated at 37 °C with 1.5 mM L-DOPA (Millipore Sigma) in the absence or presence of ampyrone (Millipore Sigma), and catalytic activities were monitored over 10 h by measuring dopachrome formation at 475 nm (ε_dopachrome_ = 3700 M^-1^ cm^-1^). For the measurement of hTYR^WT^ activity during unfolding, the enzyme was incubated with urea at concentrations ranging from 0 to 8 M for 1 hour, and activity was determined using 1.5 mM L-DOPA as a substrate by measuring dopachrome formation at 475 nm in absence or presence of 5 mM ampyrone.

To evaluate the potency of additional small-molecule modulators, hTYR^WT^ was incubated with selected potential inhibitors at compound-specific concentrations ranges, selected based on preliminary activity screening. Enzymatic activity was measured at 37 °C by recording absorbance at 475 nm every 1 minute for 120 minutes. Representative time curves were generated for each compound to monitor kinetic effects. For IC₅₀ determination, endpoint absorbance values at 120 minutes were used to calculate residual enzymatic activity across the tested concentration range. Concentration–response curves were fitted using a four-parameter nonlinear regression model in GraphPad Prism, version 10.4.0 (GraphPad Software, LLC), and IC₅₀ values were calculated accordingly.

***Michaelis-Menten kinetics***

The diphenol oxidase and monophenolase reaction rates (V) of hTYR^WT^ and hTYR^P406L^ in the presence or absence of ampyrone were determined using L-DOPA or L-tyrosine as a substrate at concentrations ranging from 0.098 to 6 mM or 0.023 to 0.75 mM, respectively. All assays were performed at 37 °C in 10 mM sodium phosphate buffer, pH 7.4. Absorbance was measured at 475 nm using the SpectraMax i3 multi-mode detection platform (Molecular Devices). The Michaelis-Menten constant (*K*_m_) and maximal velocity (V_max_) were calculated from Michaelis-Menten plots using GraphPad Prism, version 10.4.0 software. The enzyme turnover rate, k_cat_, was determined as V_max_/E_t_, where E_t_ represents the protein concentration (10 nM).

***Computational analysis***

*Model Preparation & Molecular Dynamics*. A glycosylated homology model of hTYR^WT^ was previously generated using the NEI Data Commons Ocular Proteomes TYRP1 atomic model (PDB:5M8L, <https://neicommons.nei.nih.gov/#/proteomeData>) [4]. The hTYR^P406L^ variant was created from this model using the Edit > Swap > Residue function in YASARA Structure, version 25.1.13. Both the hTYR^WT^ and hTYR^P406L^ models were energy minimized using the YASARA Options > Choose experiment > Energy minimization function. The 3D conformer structure of ampyrone was obtained from the PubChem compound database in SDF format and then converted to PDB format in UCSF Chimera, version 1.19. For the ampyrone docking experiments, which were carried out using the VINA software implemented in YASARA Structure, ten timeframes ranging from 0 to 10 ns were selected from the molecular dynamics (MD) simulation for hTYR^WT^ and hTYR^P406L^ models. Both hTYR^WT^ and hTYR^P406L^, with and without ampyrone, underwent triplicate 100 ns MD simulations (MD_run.mcr) using the AMBER14 force field (total of 12 simulations), with snapshots saved every 1 ns. Simulations were performed at 298 K, pH 7.4, and 0.9% NaCl in a cubic cell extending 10 Å beyond the protein (94.6 Å × 94.6 Å × 94.6 Å). Initial atomic velocities were varied by altering the Randomized Seed for each simulation.

*Principal Component Analysis (PCA)*. Concatenated.xtc trajectory files were aligned to a reference structure, and alpha carbon coordinates for all 449 atoms were extracted and inputted into GraphPad Prism, version 10.4.1. PCA was performed with data standardization and parallel analysis, in accordance with Prism’s guidelines. Loadings |x| ≥ 0.7 for PC1 and PC2 subspaces were projected onto the corresponding PDB structures.

*Molecular docking.* L-tyrosine and L-DOPA were procured from PubChem and docked to 100 ns protein structures for all four protein environments in YASARA Structure using the ‘dock_run.mcr’ macro and VINA. Docking was restricted to a 20 Å^3^ box centered around the copper ions. For each ligand, 25 docking runs were performed, and the top three binding poses were selected based on orientation within 4 Å of the copper atoms and hydroxyl groups oriented towards the active site.

*Visualization, RMSD, RMSF, and SASA calculations.* Structural alignments and visualizations were performed in UCSF Chimera, version 1.18.0. The superposition of hTYR^WT^ and hTYR^P406L^ models after MD simulation used the Tools > Structure Comparison > Matchmaker tool. Distances between atoms were calculated across trajectories using the Graphics > Labels > Bonds > Graph > Save tool in the Visual Molecular Dynamics program, VMD, version 2.0.0. The root-mean square deviations (RMSD) and the root mean-square fluctuations (RMSF) for each trajectory were calculated in VMD by aligning frames to the 0 ns protein structure using the Trajectory > Align. The RMSF values were averaged using a TCL file in VMD’s TK Console. RMSD was calculated using the RMSD-Trajectory > Align > RMSD. Solvent-accessible surface area (SASA) for the active site was calculated in YASARA using the Analyze > Surface Area of > Object for the 100 ns protein structure for hTYR^WT^ and hTYR^P406L^ with and without ampyrone. ΔSASA was determined as the difference between bound and unbound forms for each protein.

*DCCM and Porcupine Dynamic analysis*. Python, version 3.13 was used to generate a dynamical cross-correlation matrix (DCCM) from aligned alpha carbon coordinates. The correlation matrix was calculated as normalized dot products (range -1 to +1). A 2D porcupine plot was created using principal component eigenvectors from GraphPad Prism, version 10.4.1 to the direction and magnitude of each vector for select active site residues.

*Free Energy Landscape*. Free energy landscapes (FEL) were created using the Grossman weighted histogram analysis method (WHAM) for each protein with and without ampyrone to compare the highest probability conformations of the binding site. Reaction coordinates were defined by distances between gate-keeping residues: K334, F347, and V377, calculated using VMD Graphics > Labels > Bonds > Graph > Save. WHAM was run with 21 bins, tolerance of 10^−5^, 0 spring constant, 298 K temperature, and no periodic boundary padding. The minimum and maximum bins were determined by the corresponding values for each of the data sets.

***Mouse and human melanocyte culture***

Mouse melanocytes (melan-ink4a^-/-^) were obtained from the Wellcome Trust Functional Genomics Cell Bank (St. George’s, University of London, UK), tested, authenticated, and cultured at 37 °C and 10% CO_2_ in a Thermo Scientific Heracell Vios 160i incubator in Roswell Park Memorial Institute (RPMI) 1640 supplemented with 10% fetal bovine serum (FBS), 1% penicillin-streptomycin, 2 mM glutamine, 200 nM TPA, and 200 pM cholera toxin at 10% CO_2_. NBMEL 1284 and C4 normal melanocytes and OCA1B-1235 (TYR c.649C>T and c.[301=;575C>A;1205G>A]) and OCA1B-1125 (TYR c.242C>T and c.[301=;575C>A;1205G>A]) human melanocytes were obtained from the Biospecimen Core of the Yale Specialized Programs of Research Excellence (SPORE) in Skin Cancer (New Haven), Harvard Medical School, and the National Institutes of Health, respectively. Cells were cultured at 37 °C and 5% CO_2_ in a Thermo Scientific Heracell Vios 160i incubator in Ham’s F-10 (Gibco) supplemented with 5% fetal bovine serum (Corning), 2 mM L-Glutamine (Gibco), 250 ng/mL Amphotericin B (Gibco), 1% Penicillin Streptomycin (100 Units/mL and 100 μg/mL, respectively) (Gibco), 5 ng/mL basic fibroblast growth factor (Peprotech), 10 ng/mL Endothelin 1 (Sigma-Aldrich), 7.5 μg/mL 3-isobutyl-1-methylxanthine (Sigma-Aldrich), 30 ng/mL cholera toxin (Sigma-Aldrich), and 3.3 ng/mL Phorbol-12-myristate-13-acetate (Fisher Scientific). Media was replaced every third day. Mycoplasma testing was performed quarterly using commercial kits.

***LC-MS tyrosine tracing***

Mouse melanocytes (ink4a^-/-^) were plated in 6-well plates and grown to approximately 80-90% confluence, then incubated in [U-^13^C] tyrosine medium containing vehicle or ampyrone (2 mM) for 20, 40, and 60 minutes at 10% CO_2_. Cells were quickly washed twice with ice-cold PBS, followed by a quick ddH_2_O rinse and metabolite extraction using -70 °C 80:20 methanol: water (LC-MS grade methanol, Fisher Scientific). The cell-methanol mixture was subjected to bead-beating for 45 sec using a Tissuelyser cell disrupter (Qiagen). Extracts were centrifuged for 10 min at 13.2k rpm to pellet insoluble material and supernatants were transferred to clean tubes. The extraction procedure was repeated two additional times, and all three supernatants were pooled, dried in a speed-vac (Savant) and stored at -80 °C until analysis. The methanol-insoluble protein pellet was solubilized in 0.2 M NaOH at 95 °C for 20 min and quantified using the BioRad DC assay. On the day of metabolite analysis, dried cell extracts were reconstituted in 70% acetonitrile at a relative protein concentration of 2.5 mg/mL, and 8 µL of this reconstituted extract was injected for LC-MS-based targeted stable isotope profiling. Cell extracts were analyzed by LC-MS as described previously [5-9] using a platform comprised of an Agilent Model 1290 Infinity II liquid chromatography system coupled to an Agilent 6550 iFunnel time-of-flight MS analyzer. Chromatography of metabolites utilized aqueous normal phase (ANP) chromatography on a Diamond Hydride column (Microsolv). Mobile phases consisted of: (A) 50% isopropanol, containing 0.025% acetic acid, and (B) 90% acetonitrile containing 5 mM ammonium acetate. To eliminate the interference of metal ions on chromatographic peak integrity and electrospray ionization, Ethylenediaminetetraacetic acid (EDTA) was added to the mobile phase at a final concentration of 6 µM. The following gradient was applied: 0-1.0 min, 0% B; 1.0-15.0 min, to 20% B; 15.0 to 29.0 min, 50% B; 29.1 to 37 min, 99% B. Raw data were analyzed using MassHunter Profinder 8.0 and MassProfiler Professional (MPP) 14.9.1 software (Agilent technologies). An in-house untargeted stable isotope tracing (USIT) workflow [9, 10] was employed to obtain all possible fates of [U-^13^C] tyrosine and quantitative information on the relative incorporation of tyrosine–derived metabolites based on the stable isotope labeling pattern. USIT used the Agilent untargeted metabolite profiling software [MassHunter Qualitative Analysis 8.0, MassProfinder 8.0 and MassProfiler Professional (MPP 14.9)] for an initial targeted identification of differentially expressed metabolites in cells grown in [U-^13^C] tyrosine-supplemented media.

***Measurement of melanin content by flow cytometry***

NBMEL 1284 and C4 normal human melanocytes and OCA1B-1235 and OCA1B-1125 human melanocytes were treated with normal culture media in 5% CO_2_ and either control or with 0.2 mM ampyrone for 48 hours. Following treatment, cells were trypsinized and resuspended in 100 µL PBS without calcium and magnesium (Corning). Flow cytometry was performed using a BD Fortessa X20 5 laser analyzer. Data collection included forward scatter, side scatter, and scatter intensities at 355 nm. A minimum of 2000 events were collected and analyzed per replicate sample. All analysis was performed using FCExpress (DeNovo Software). Events were gated by side scatter and forward scatter to isolate live cells prior to further processing. Differential light scattering due to melanin was captured as mean scatter intensity through a 379/28 nm bandpass filter using a 355 nm ultraviolet laser. Mean scatter was statistically analyzed using unpaired Student’s t-test comparing vehicle to ampyrone treated cells.

***In vitro 3D tissue culture, treatment and imaging***

MelanoDerm™ (MEL-312-B; MatTek Corp.) was used as a human skin epidermis model. This 3D human skin equivalent was derived from Black donors and contains normal melanocytes and keratinocytes. MelanoDerm™ was grown at the air-liquid interface in EPI-100-NMM-3 medium provided by the manufacturer. The cells used to obtain the 3D model were co-cultured on a collagen-coated membrane to form a multilayered, highly differentiated model of the human epidermis. The apical surfaces of the reconstructed tissues were exposed to air, while the bottom surfaces remained in contact with the culture medium [11]. Prior to ampyrone treatment, tissues were washed with 1 mL PBS. Ampyrone was dissolved in EPI-100-NMM-3 medium to a final concentration of 0.2 mM. Negative control samples were treated only with medium, while the positive control samples were treated only with EPI-100–NMM-113 medium from the manufacturer. Ampyrone in medium was applied to MelanoDerm™ every other day for 21 days. The epidermal images were captured using a stereo microscope (ZEISS SteREO Discovery.V12). Melanocyte images were captured using an integrated digital inverted microscope (EVOS™ M5000 Imaging System, Thermo Fisher Scientific). On the 21^st^ day of the experiment, samples were processed, embedded in paraffin blocks, and sectioned to a thickness of 5 μm onto slides, which were stained with Fontana-Masson (FM) to highlight melanin pigmentation and hematoxylin-eosin (H&E) to stain for normal structure following standard methods (Histoserv, Inc.). Images were captured using the inverted widefield microscope (Zeiss Definite Focus). Quantitative analysis of FM-stained tissue for accumulated melanin was performed using ImageJ software.

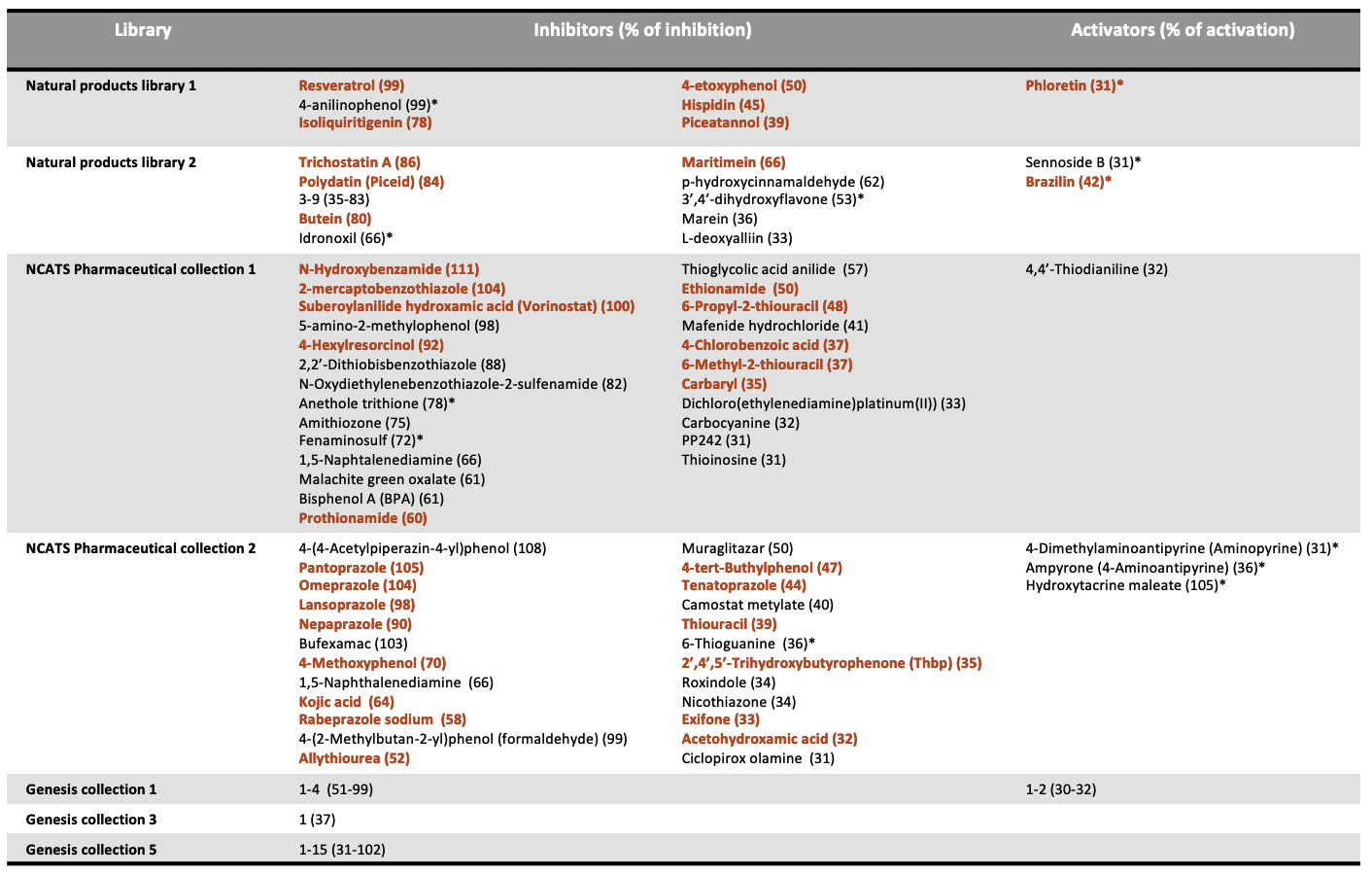

**Supplemental Table S1.** Compounds identified as inhibitors or activators of the hTYR^WT^ from a primary HTS performed with a single test concentration of 115 µM. The compounds indicated by red font were previously described as tyrosinase inhibitors. *Compounds analyzed in this study.

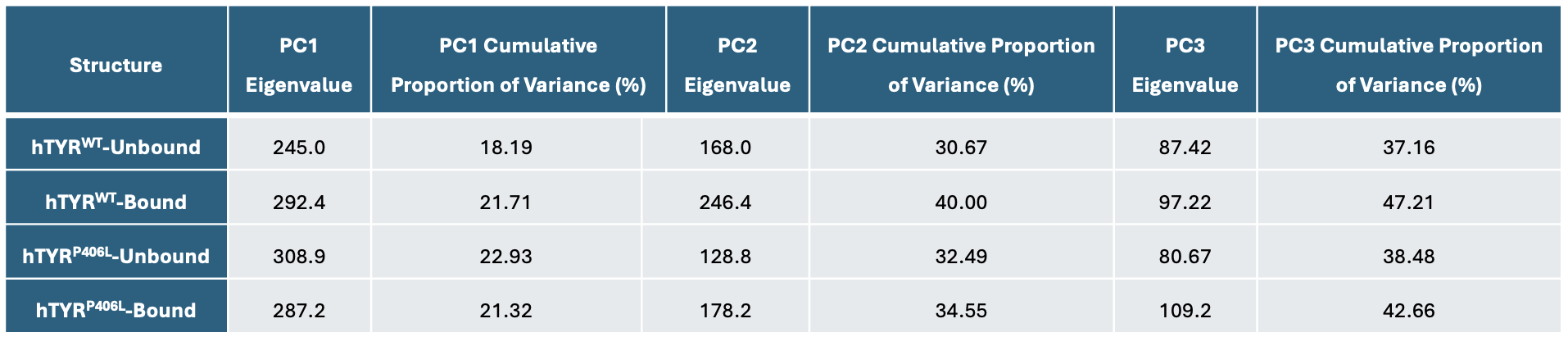

**Supplementary Table S2. Principal components summary.** This table provides specific values of variance and Eigen used in the modeling of ampyrone bound to hTYR^WT^ and hTYR^P406L^ enzymes.

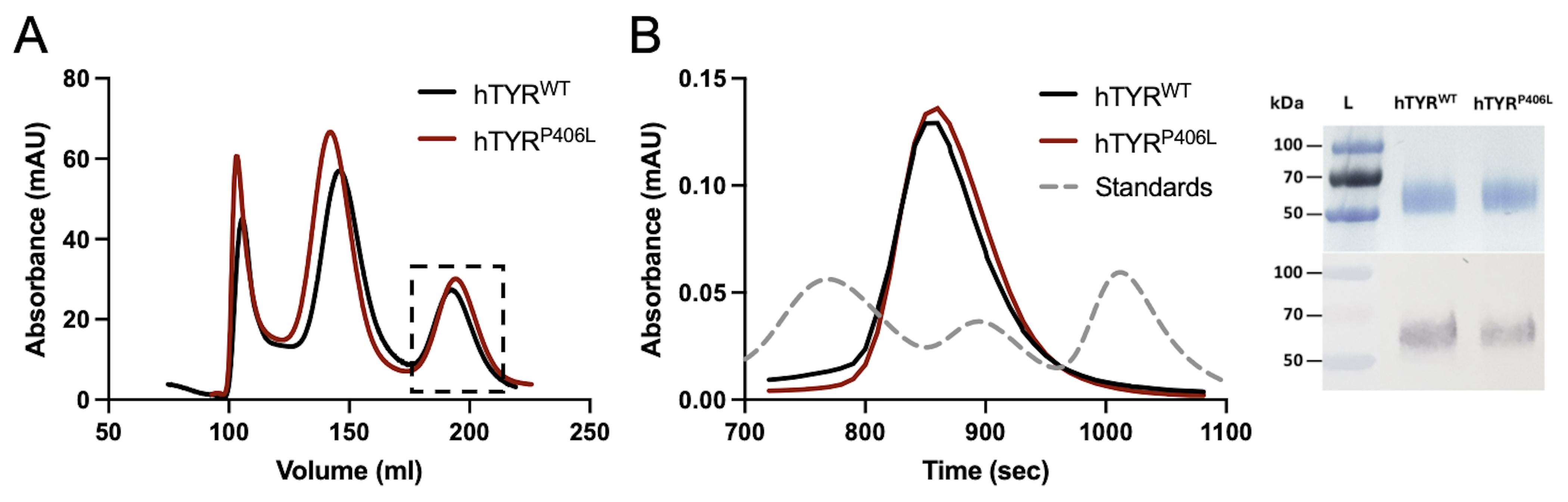

**Supplemental Figure S1. Purification of recombinant hTYR^WT^ and hTYR^P406L^ enzymes.** A) Chromatography profiles of hTYR^WT^ (black) and hTYR^P406L^ (red) eluted from the HiPrep 26/60 Sephacryl S-300 column after immobilized metal affinity chromatography (IMAC). Both proteins appear in the peak marked within the dotted frame. B) hTYR^WT^ (black) and hTYR^P406L^ (red) eluted from the Superose 12 GL 10/300 column using a Bio-Logic Duo-Flow Maximizer workstation. The grey dashed line indicates Bio-Rad molecular weight standards (gamma globulin 155.0 kDa, ovalbumin 44.0 kDa, and myoglobin 17.0 kDa). SDS-PAGE (inset, top) shows protein purity, and an anti-TYR (T311, 1:2000) Western blot (inset, bottom) confirms the molecular identity of the purified proteins. The molecular weights of the protein ladder are labeled at 50, 70, and 100 kDa.

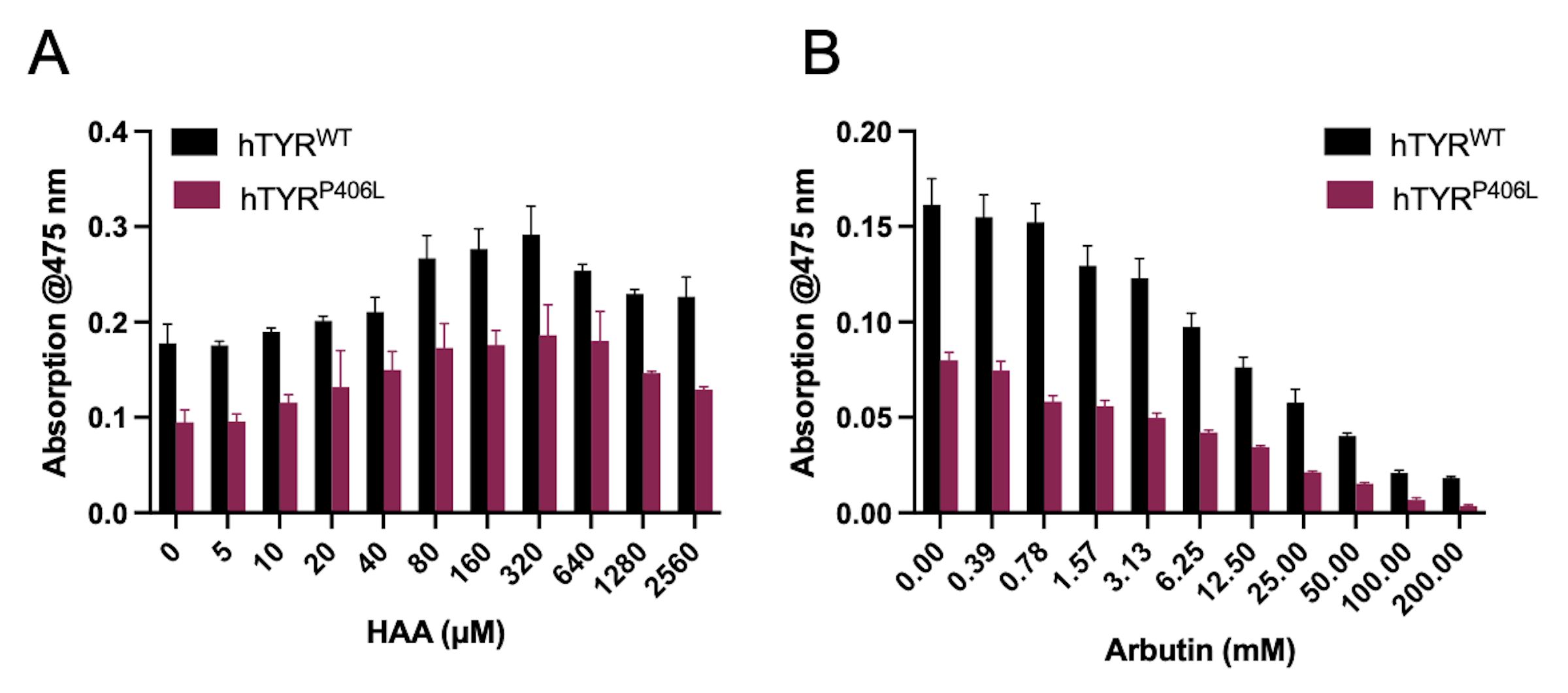

**Supplemental Figure S2. Functional validation of hTYR^WT^ and hTYR^P406L^.** Functional validation of hTYR^WT^ (black) and hTYR^P406L^ (red) with a known activator, 3-hydroxyanthranilic acid (HAA) (A), and a known inhibitor, arbutin, of *in vitro* diphenol oxidase activity (B).

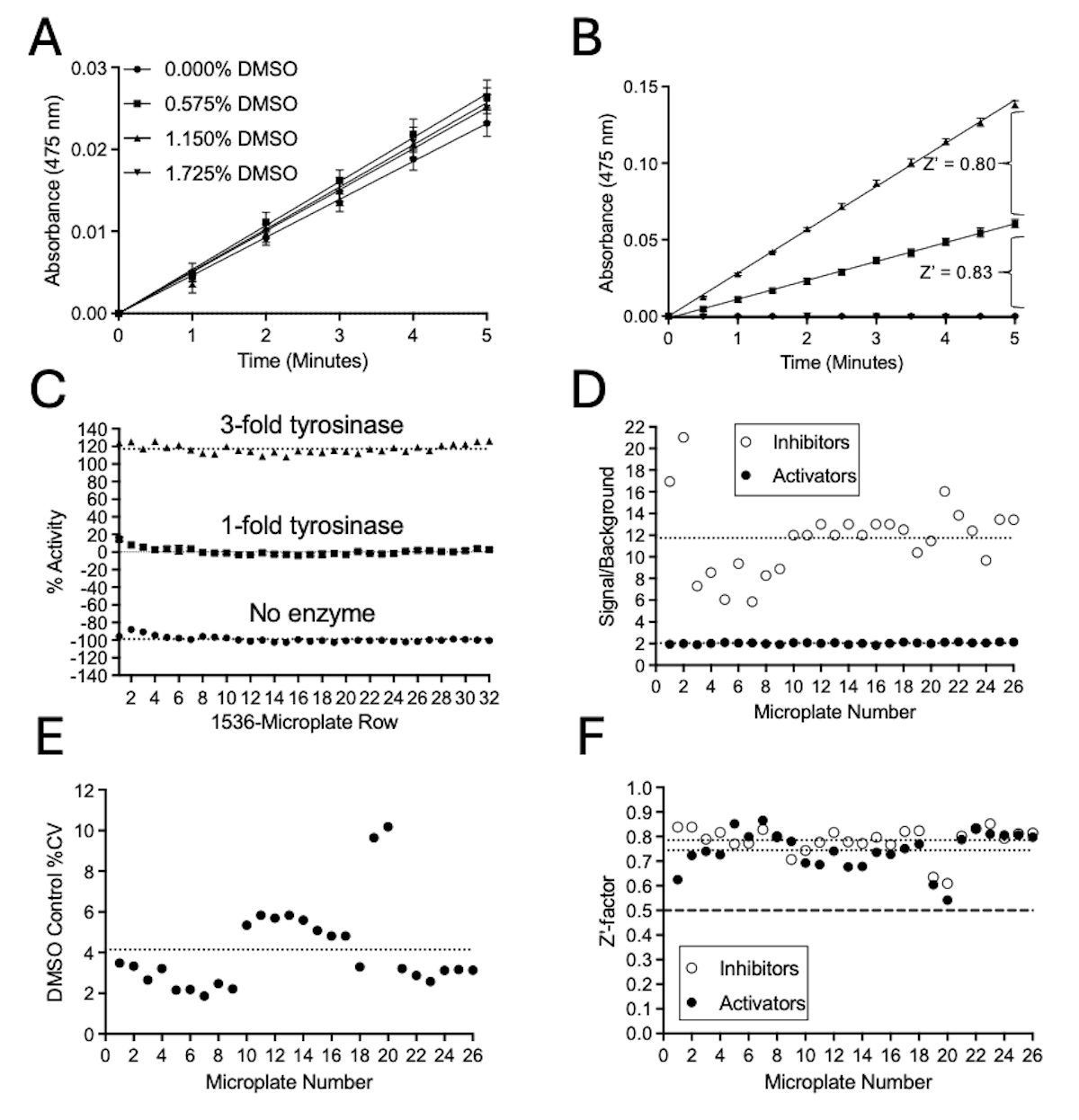

**Supplemental Figure S3. Validation and performance of a miniaturized HTS assay for simultaneously identifying both activators and inhibitors of hTYR^WT^ diphenol oxidase activity *in vitro*.** A) The optimized diphenol oxidase activity assay was performed with hTYR^WT^ (0.75 µM) in the absence and presence of increasing concentrations of DMSO. No reduction in enzymatic activity was observed at concentrations up to 1.73%. All data show the mean ± SD (*n* = 3 technical replicates). B) Kinetics of the optimized assay controls including the high activity control (triangles, 3-fold hTYR^WT^, 4.5 µM), baseline control (squares, hTYR^WT^, 1.5 µM) and low activity control (diamonds, no enzyme). Together, the controls enabled the assessment of assay performance to screen for activators (Z’ = 0.8) and inhibitors (Z’ = 0.83). All data show the mean ± SD (*n* = 3 technical replicates). C) Percent activity of controls in a 1536-well microplate including the high activity control (*n* = 32), baseline control (*n* = 32) and low activity control (*n* = 32). D) Among 26 microplates of the primary HTS, the average signal-to-background for the low activity control (no enzyme, open circles) was 11.75 (dotted line) and 2.0 for the high activity control (3-fold hTYR^WT^, filled circles). E) The average coefficient of variation of the DMSO-treated baseline control for 26 microplates was 4.1% (dotted line). F) For the low activity control (open circles), Z’-factor values ranged between 0.61 and 0.85, with an average of 0.79 (dotted line) among 26 microplates. For the high activity control (dotted line), Z’-factor values ranged between 0.54 and 0.87, with an average of 0.74 (dotted line) among 26 microplates.

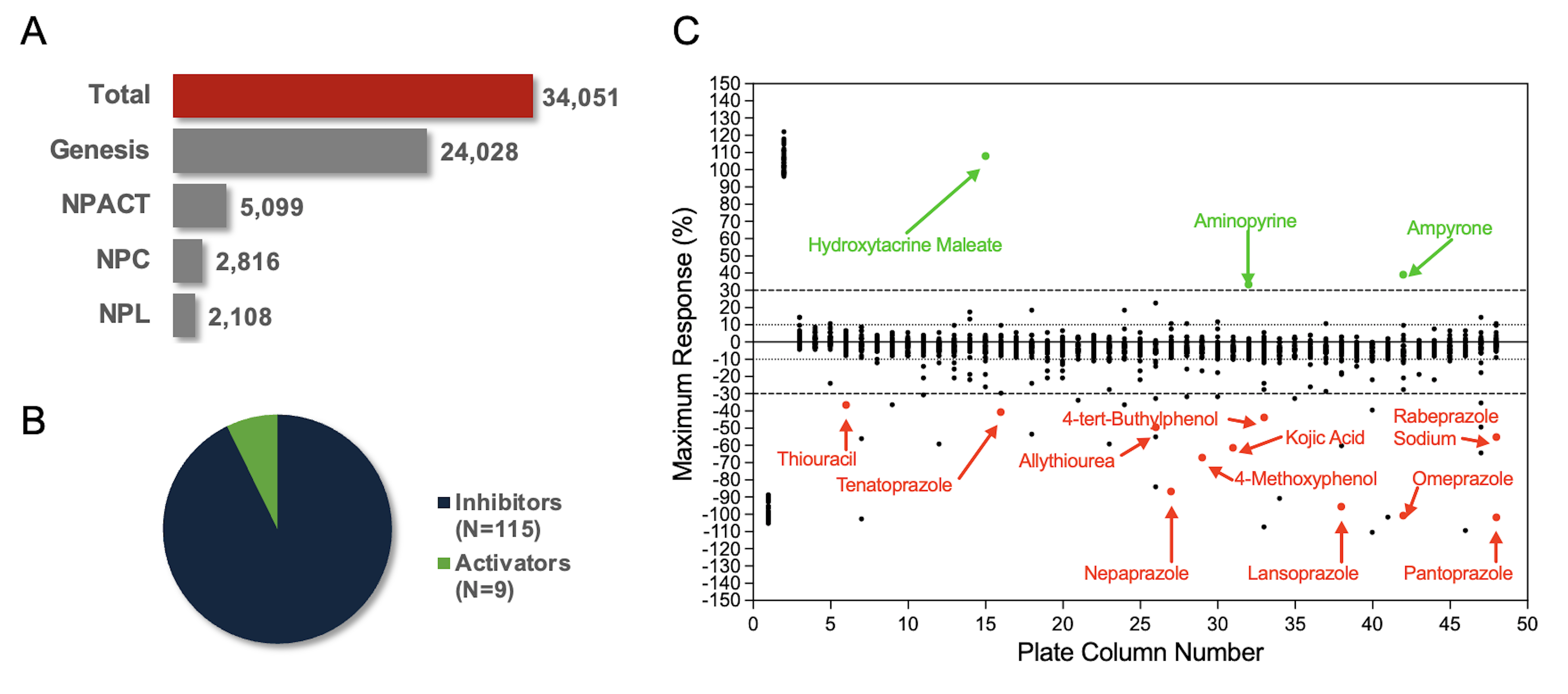

**Supplemental Figure S4. Identification of hTYR^WT^ activators and inhibitors from an HTS.** A) An HTS of hTYR^WT^ was conducted with a collection of 34,051 compounds from four chemical libraries: the Genesis Diverse Chemical Library (Genesis), the NCATS Pharmacologically Active Chemical Toolbox (NPACT), the NCATS Pharmaceutical Collection (NPC), and the Natural Products Library (NPL) at a single concentration of 115 μM. B) A total of 115 inhibitors and nine activators were identified. C) Activity data resulting from 1,408 compounds of the NCATS Pharmaceutical Collection 2 with names of activators (green) and inhibitors (red) annotated **(see Supplemental Table S1).**

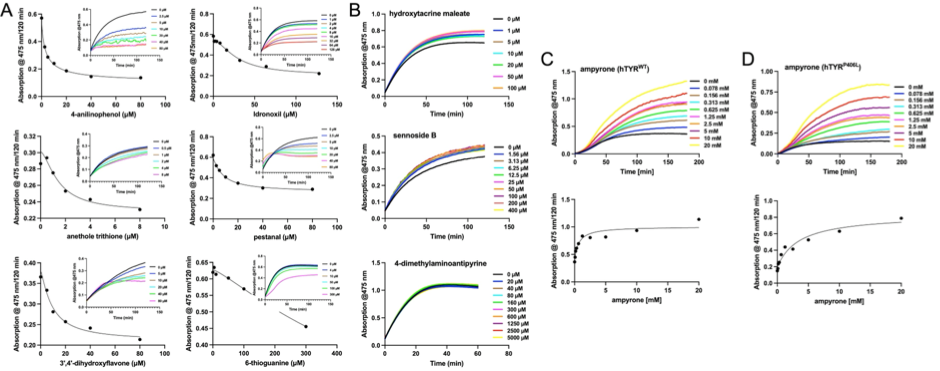

**Supplemental Figure S5. Confirmation of inhibitor and activator hits.** A) Concentration-response graphs for hTYR^WT^ diphenol oxidase activity inhibitors. Absorbance at 475 nm after 120 min was measured following incubation at 37ºC with increasing concentrations of 4-anilinophenol, idronoxil, anethole trithione, pestanal, 3’,4’-dihydroxyflavone, and 6-thioguanine. Data were fitted with a nonlinear regression model to estimate IC_50_. Insets show time-dependent inhibition profiles for each compound. B) Representative time curves of the hTYR^WT^ diphenol oxidase activity, measured by absorbance at 475 nm, in the presence of the potential activators: hydroxytacrine maleate, sennoside B, and 4-dimethylaminoantipyrine at the indicated concentrations. C) hTYR^WT^ diphenol oxidase activity. Top: representative time curves in the presence of ampyrone. Bottom: concentration-response curve of ampyrone, witch absorbance at 475 nm after 120 min following incubation at 37ºC with increasing concentrations. Data were fitted with a nonlinear regression model to estimate EC_50_. D) hTYR^P406L^ diphenol oxidase activity. Top: representative time curves in the presence of ampyrone. Bottom: concentration-response curve of ampyrone, witch absorbance at 475 nm after 120 min following incubation at 37ºC with increasing concentrations. Data were fitted with a nonlinear regression model to estimate EC_50_. enzymes.

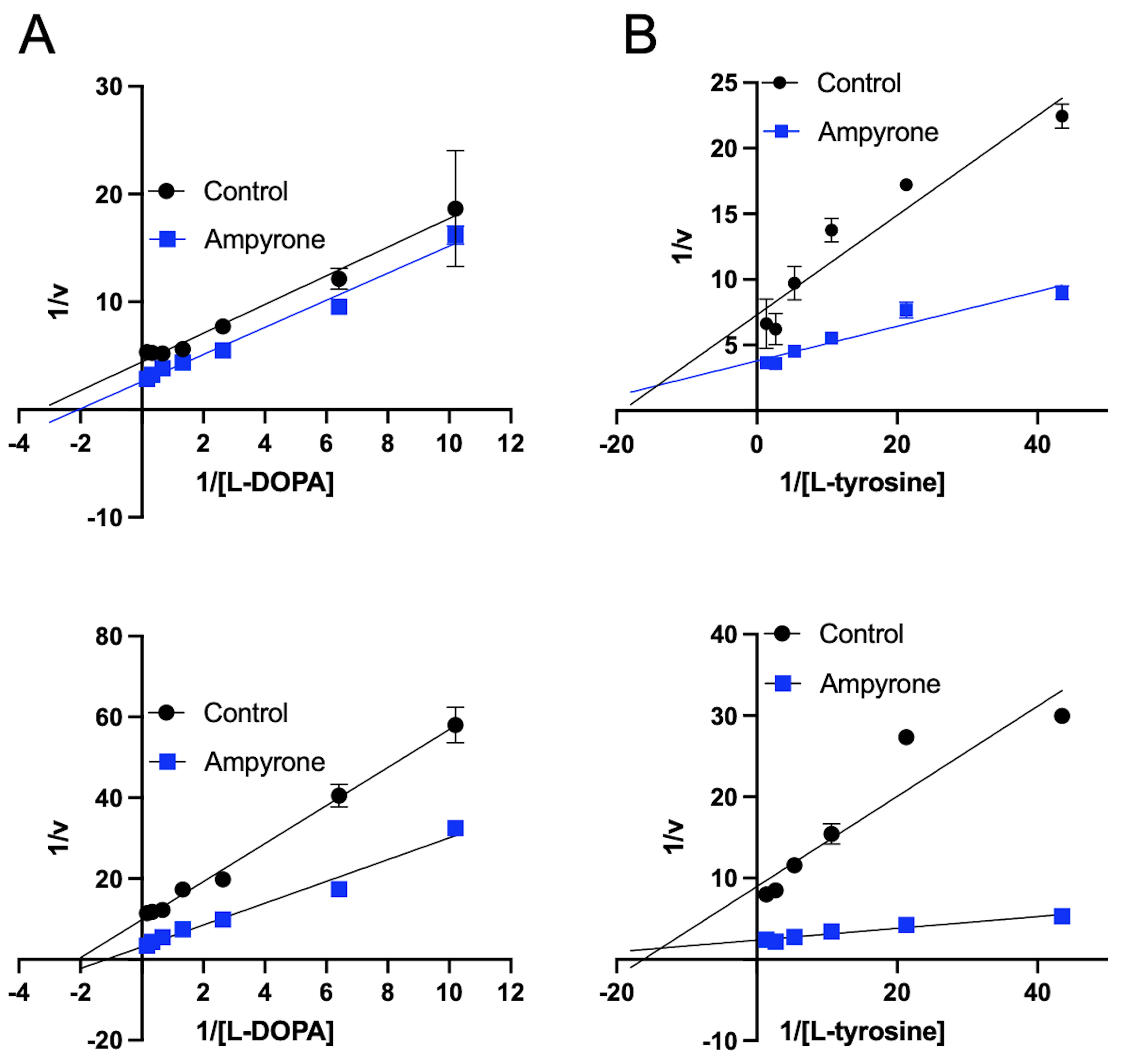

**Supplemental Figure S6. Lineweaver-Burk analysis of ampyrone’s effect on the diphenol oxidase and monophenolase activity of hTYR^WT^ and hTYR^P406L^.** Lineweaver–Burk plots of diphenol oxidase (A) and monophenolase (B) activity of hTYR^WT^ (top) and hTYR^P406L^ (bottom) enzymes measured at 37°C in the presence (ampyrone, blue) or absence (control, black) of 10 mM ampyrone, using increasing concentrations of L-DOPA or L-tyrosine, respectively. Data points were fit to the linearized Lineweaver–Burk equation in GraphPad Prism 10. Data represent the mean ± SD from three replicate experiments.

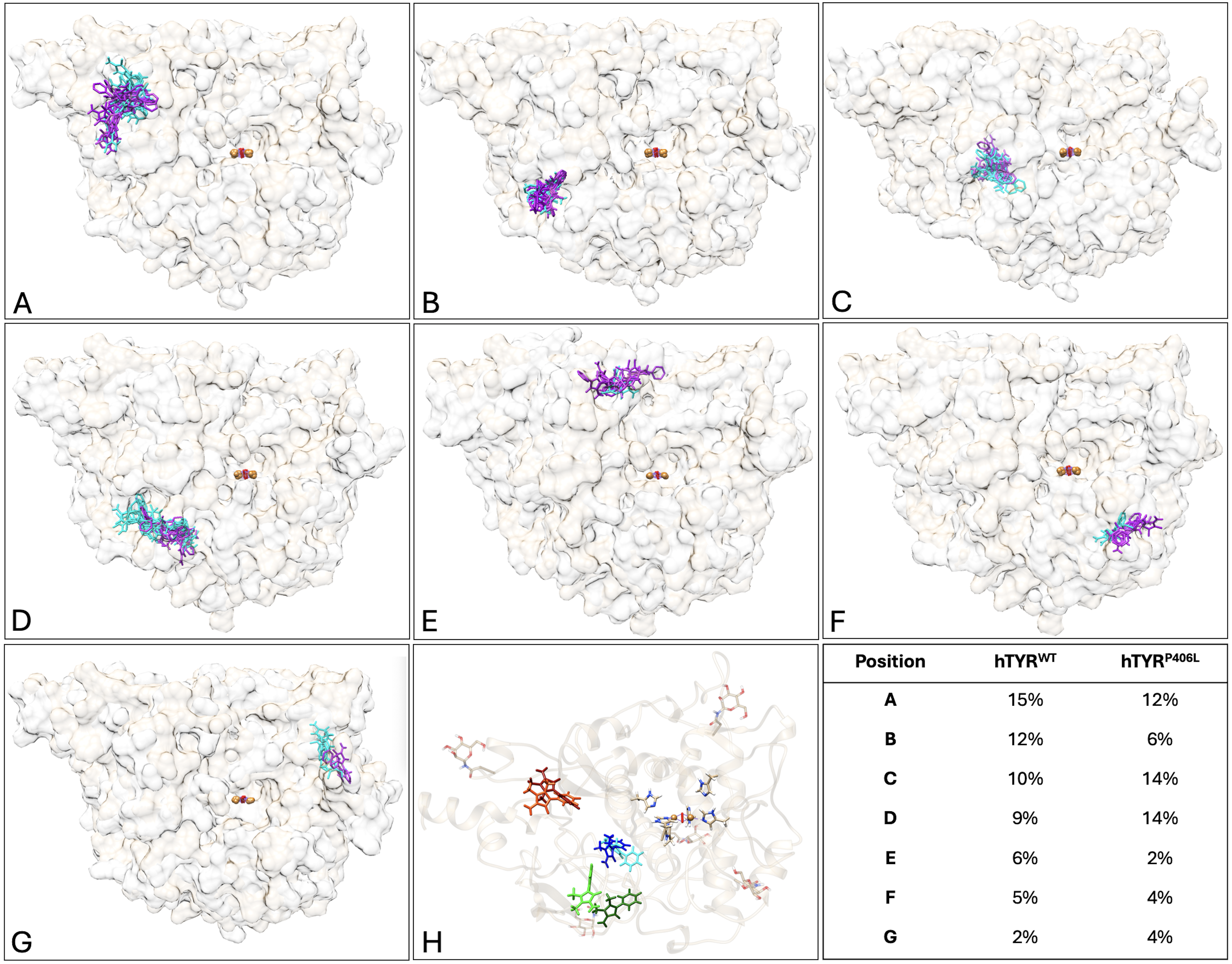

**Supplemental Figure S7. The most frequently occupied ampyrone binding positions outside the active site are similar for both hTYR^WT^ and hTYR^P406L^, as determined by molecular docking.** Ampyrone docking poses A - G) are shown in purple for hTYR^WT^ and cyan for the hTYR^P406L^. Light tan represents the surface of the superimposed hTYR^WT^, while light grey indicates the hTYR^P406L^. The copper atoms in the active sites are depicted as orange spheres. H) shows the top three binding sites for ampyrone in hTYR^WT^ (Position A/Orange, Position B/Light Green, and Position C/Cyan) and hTYR^P406L^ (Position A/Red, Position C/Royal Blue, Position D/Dark Green). The table summarizes the binding positions and their occupancy frequency.

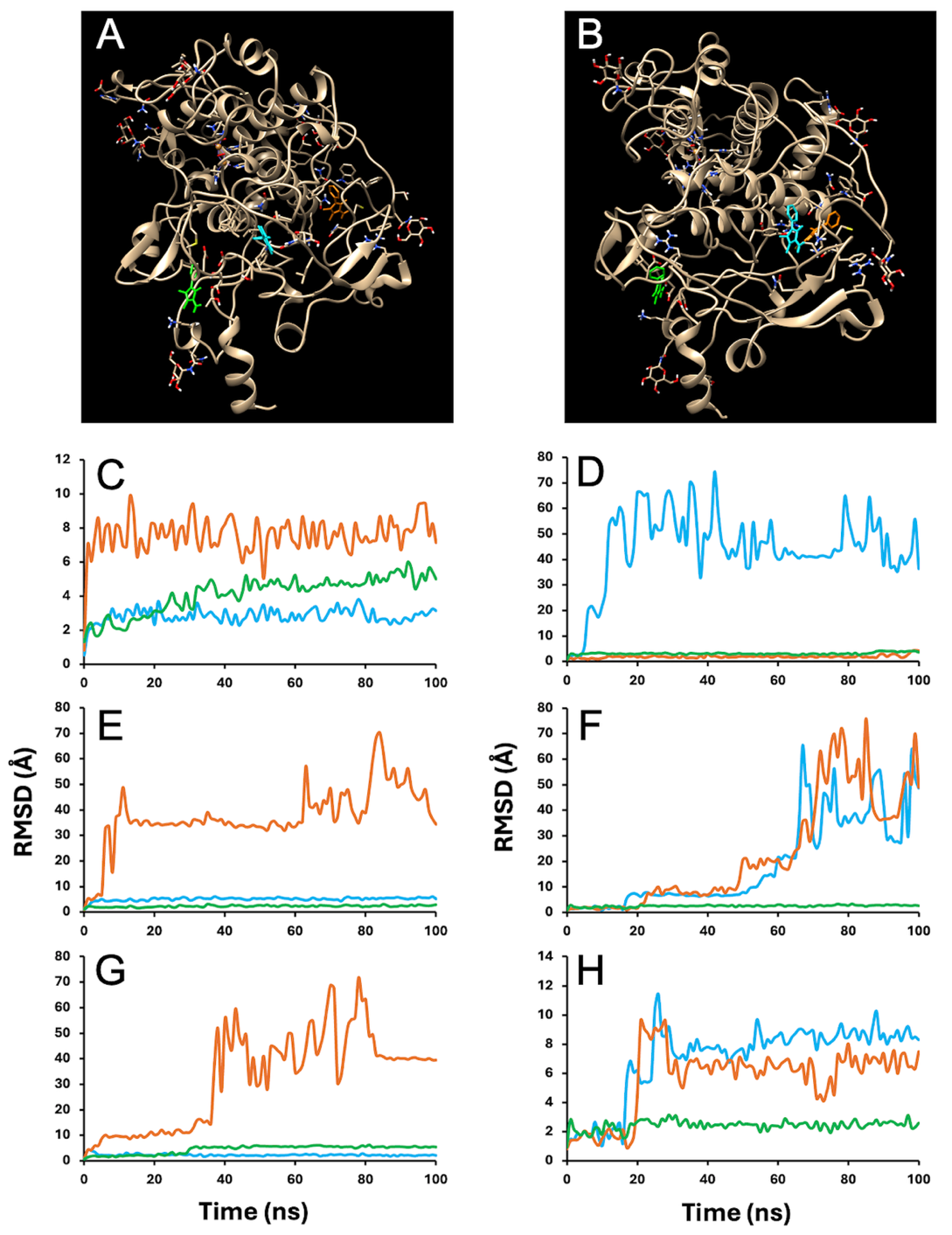

**Supplemental Figure S8. Ampyrone binding sites and stability across simulations.** Initial locations of ampyrone binding sites in hTYR^WT^ (A) and hTYR^P406L^ (B). Three ampyrone molecules are shown in blue (A1) orange (A2), and green (A3). Panels C-H: RMSD profiles for ampyrone molecules bound to hTYR^WT^ (C-E) and the hTYR^P406L^ (F-H) across three independent simulation samples. Three ampyrone molecules were bound to hTYR^WT^ and hTYR^P406L^: A1 (blue), A2 (orange), and A3 (green).

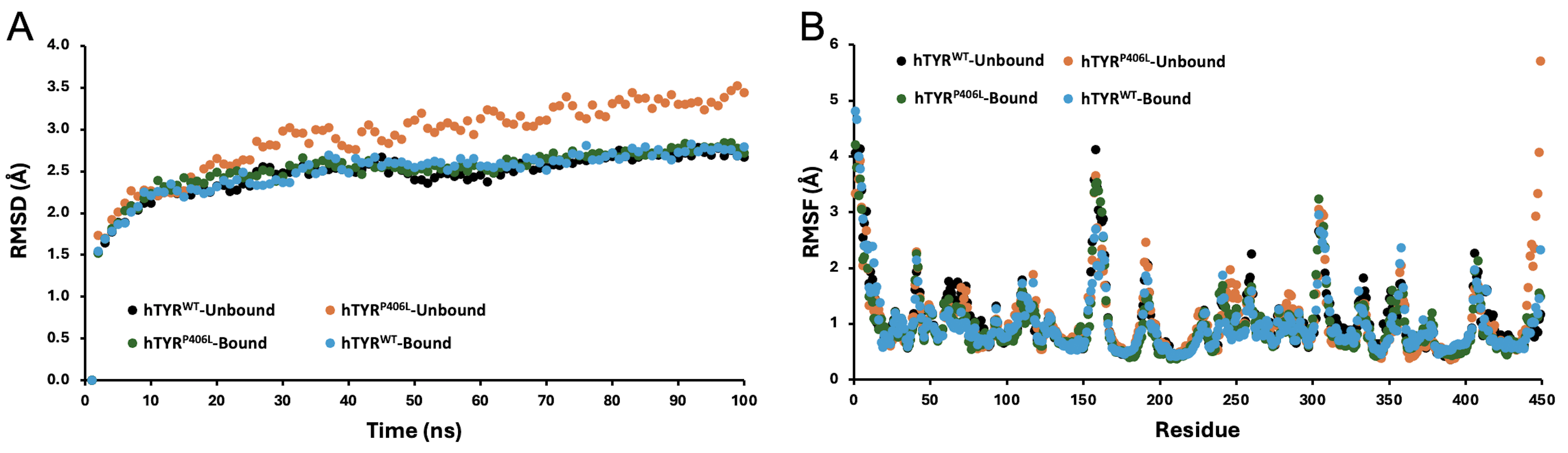

**Supplemental Figure S9. Time changes for RMSD and RMSF.** RMSD (A) and RMSF (B) for hTYR^WT^-Unbound (black), hTYR^P406L^-Unbound (orange), hTYR^P406L^-Bound (green), hTYR^WT^-Bound (blue).

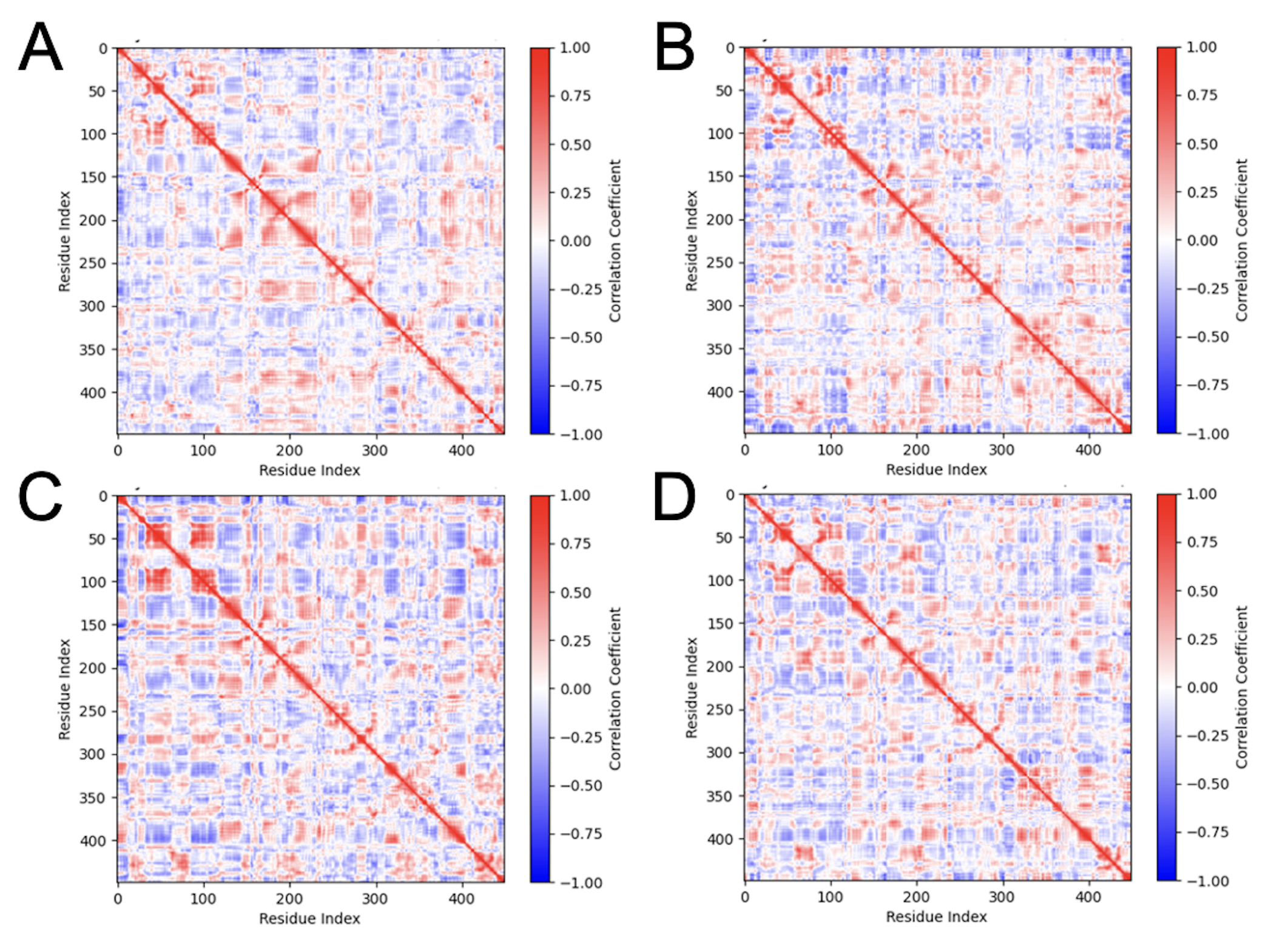

**Supplemental Figure S10. Dynamical cross-correlation matrices (DCCM) for hTYR^WT^ and hTYR^P406L^.** DCCM for hTYR^WT^ without ampyrone (A), hTYR^P406L^ without ampyrone (B), hTYR^WT^ with ampyrone (C), and hTYR^P406L^ with ampyrone.

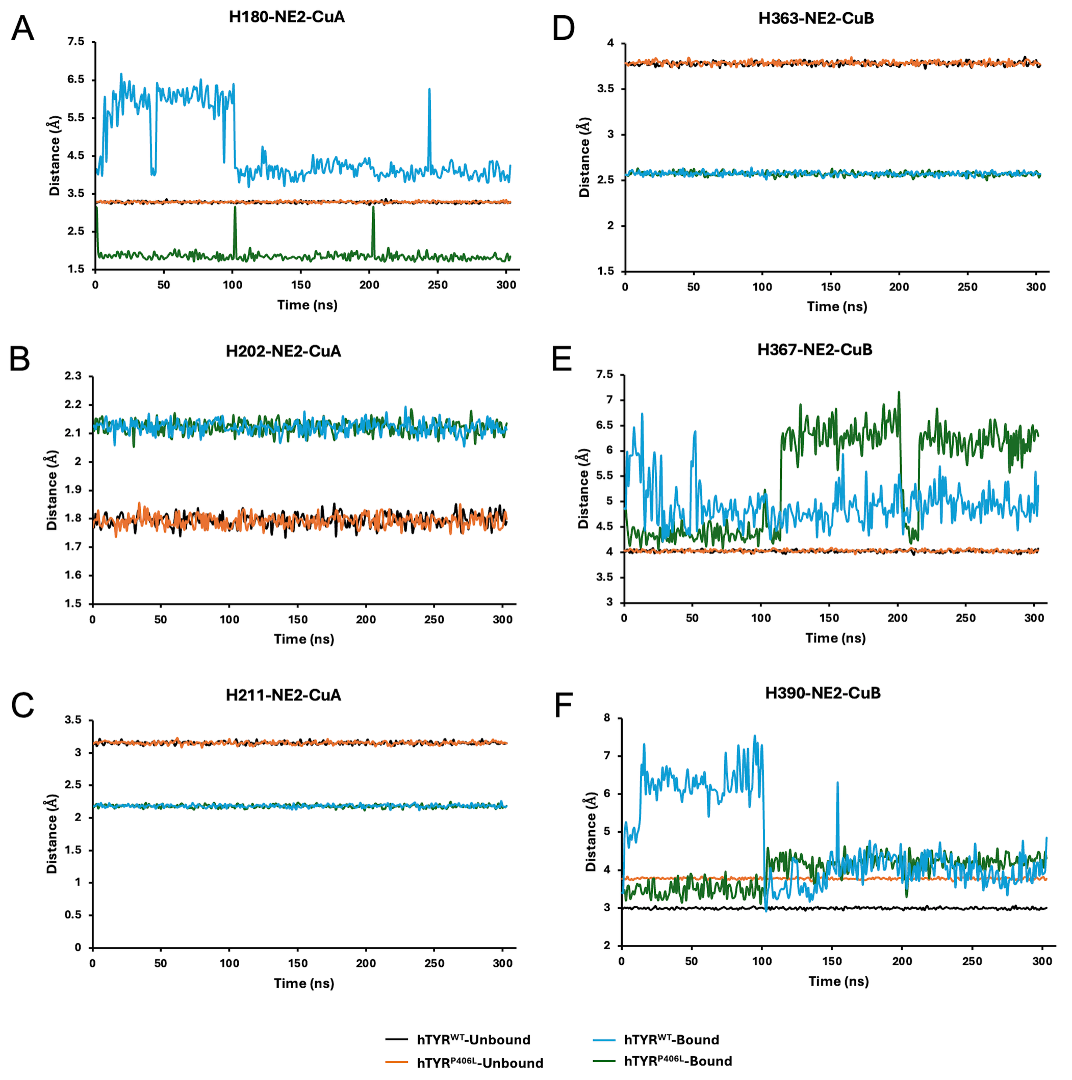

**Supplemental Figure S11. Distances between six histidines residues coordinating active site coppers (CuA and CuB)**. For each protein 2 states are considered, ‘Bound’ (ampyrone is bound to hTYR) and ‘Unbound’ (no ampyrone binding). The graphs show the distances between the CuA and NE2 atom of H180 (A), H202 (B), and H211 (C), and CuB and NE2 atom of H363 (D), H367 (E), and H390 (F) for hTYR^WT^-Unbound (black), hTYR^P406L^-Unbound (orange), hTYR^WT^-Bound (blue), hTYR^P406L^-Bound (green).

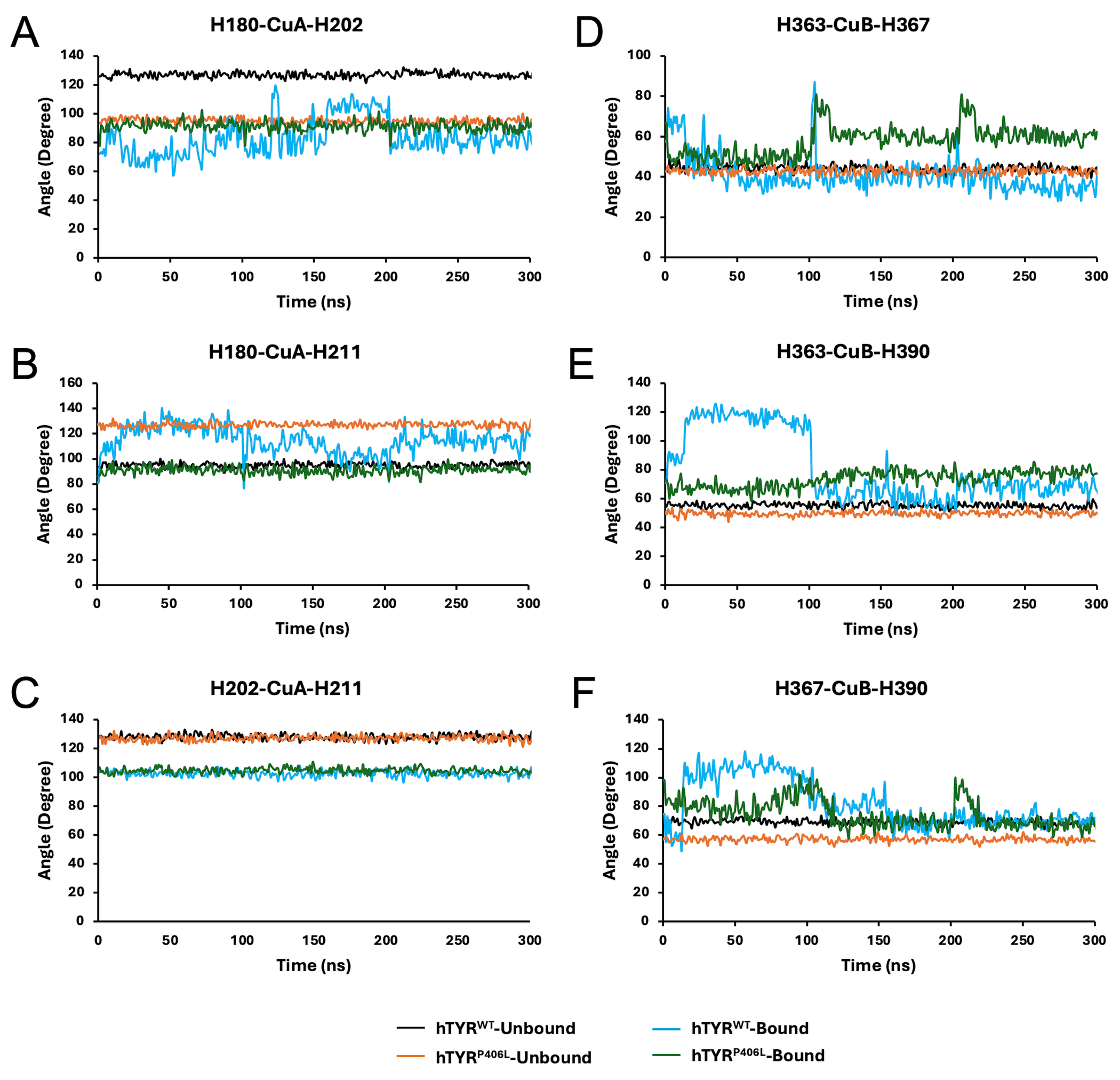

**Supplemental Figure S12. Angles of coppers coordination for six histidine residues in the TYR’s active site.** The graphs show the coordination angles between the CuA and H180/H202 (A), H180/H211 (B), H202/H211 (C), and between CuB and H363/H367 (D), H363/H390 (E), and H367/H390 (F) for hTYR^WT^-Unbound (black), hTYR^P406L^-Unbound (orange), hTYR^WT^-Bound (blue), hTYR^P406L^-Bound (green).

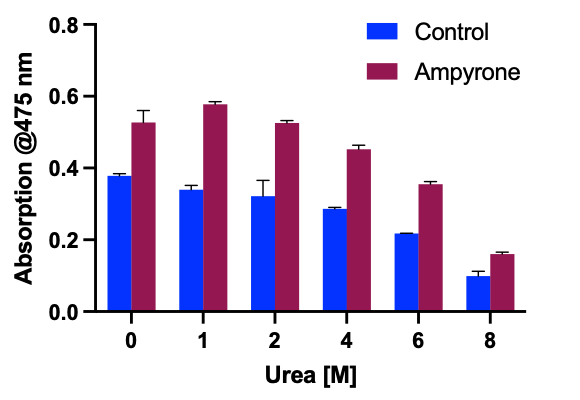

**Supplemental Figure S13. Protective effect of ampyrone on hTYR stability.** Diphenol oxidase activity of hTYR^WT^ (control, blue bars) and with 5 mM ampyrone (red bars) measured as absorbance at 475 nm after a 1hour incubation with urea at concentrations ranging from 0 to 8 M. Error bars represent standard deviations from three replicate experiments. Absorbance of the blank (samples without enzyme) was subtracted from all values.

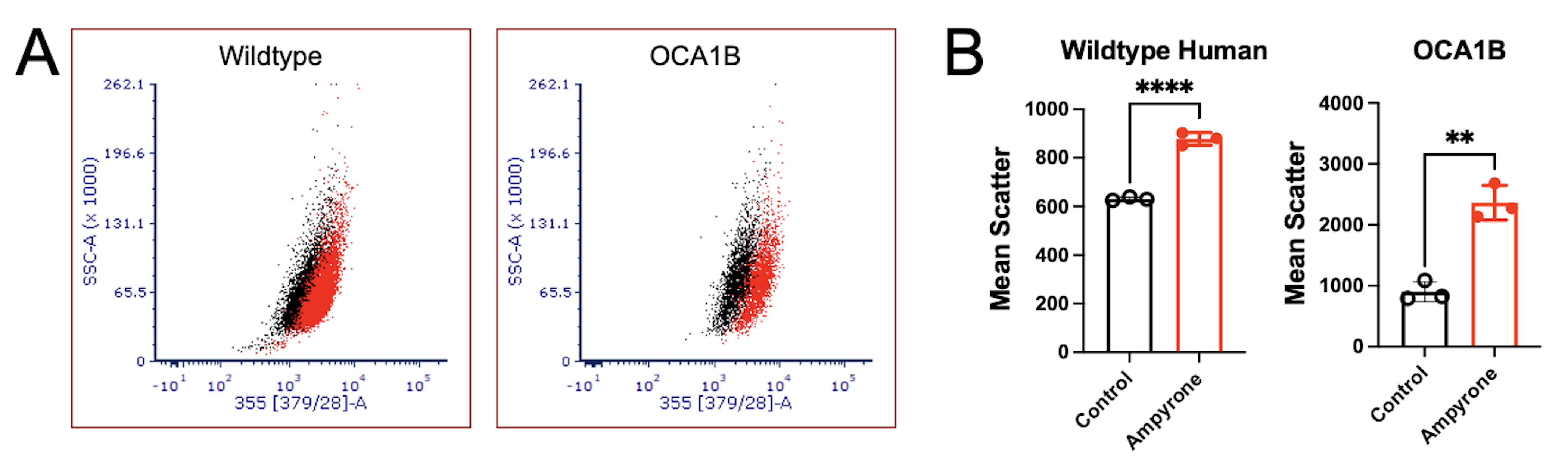

**Supplemental Figure S14.** **Ampyrone induces melanin synthesis in melanocytes.** A) Scatter plot of human wild-type (C4) and OCA1B -1125 melanocytes (see Figure 3) after three days of treatment with control (black) and ampyrone (red) showing side scatter on the Y axis and scatter at 355nm on X axis. B) Mean scatter at 355 nm for wild-type human (NB1284) and OCA1B -1235 human melanocytes after treatment with vehicle (control, black) and ampyrone (0.2 M, red) for three days. Student’s t-test, unpaired N ≥ 3 (**, P<0.01 ****, P<0.0001).

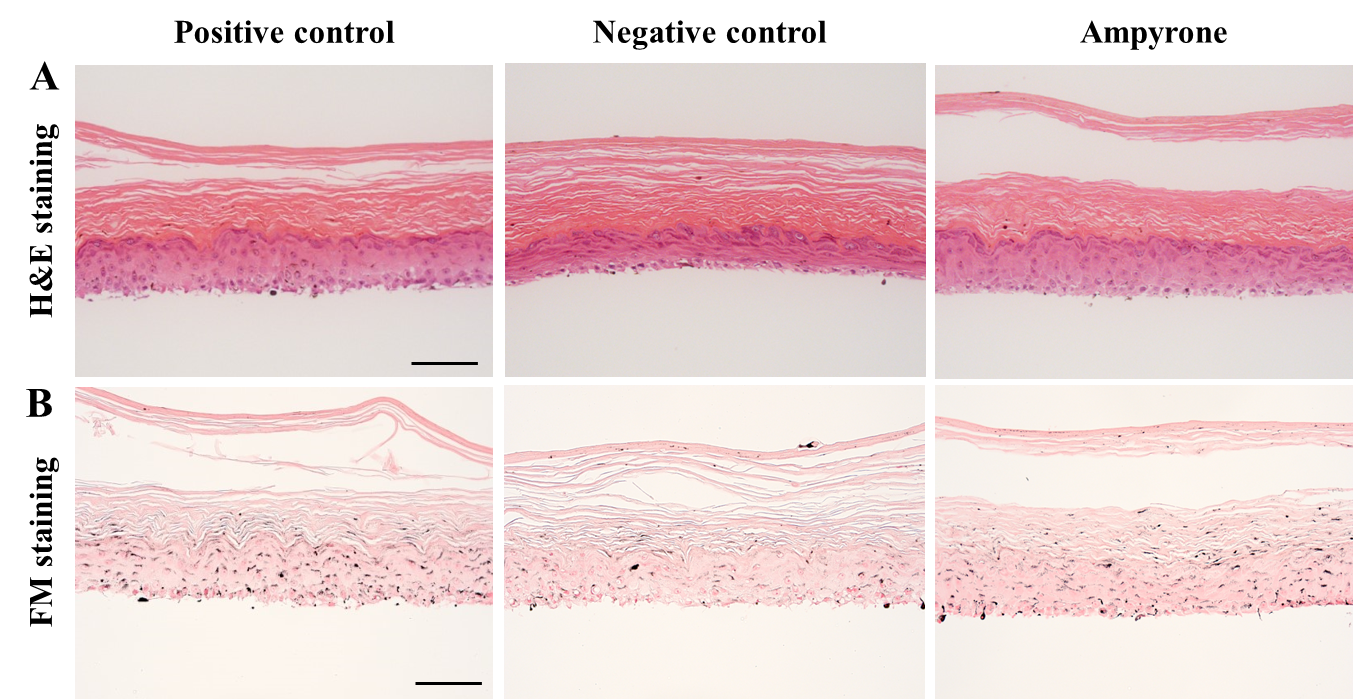

**Supplemental Figure S15. Histologic analysis of ampyrone effects on the MelanoDerm^TM^ 3D culture system**. A-B) Histological analyses (inverted wide field microscope, 20X) using hematoxylin and eosin (H&E) and Fontana-Masson (FM) staining to examine the effects of amypyone on epidermal maturation and melanin synthesis, respectively. Scale bar 100 µm.

**References**

1.	Dolinska, M.B., et al., *Albinism-causing mutations in recombinant human tyrosinase alter intrinsic enzymatic activity.* PloS one, 2014. **9**(1): p. e84494.

2.	Dolinska, M.B., et al., *Oculocutaneous albinism type 1: link between mutations, tyrosinase conformational stability, and enzymatic activity.* Pigment Cell Melanoma Res, 2017. **30**(1): p. 41–52.

3.	Dolinska, M.B., P.T. Wingfield, and Y.V. Sergeev, *Purification of Recombinant Human Tyrosinase from Insect Larvae Infected with the Baculovirus Vector.* Curr Protoc Protein Sci, 2017. **89**: p. 6 15 1–6 15 12.

4.	Patel, M. and Y. Sergeev, *Functional in silico analysis of human tyrosinase and OCA1 associated mutations.* J Anal Pharm Res, 2020. **9**(3): p. 81–89.

5.	Zhou, D., et al., *Two-pore channel 2 is required for soluble adenylyl cyclase-dependent regulation of melanosomal pH and melanin synthesis.* Pigment Cell Melanoma Res, 2024. **37**(5): p. 656–666.

6.	Chen, Q., et al., *Measurement of Melanin Metabolism in Live Cells by [U-(13)C]-L-Tyrosine Fate Tracing Using Liquid Chromatography-Mass Spectrometry.* J Invest Dermatol, 2021. **141**(7): p. 1810–1818 e6.

7.	Chen, Q., et al., *Serum Metabolite Biomarkers Discriminate Healthy Smokers from COPD Smokers.* PLoS One, 2015. **10**(12): p. e0143937.

8.	Chen, Q., et al., *Untargeted plasma metabolite profiling reveals the broad systemic consequences of xanthine oxidoreductase inactivation in mice.* PLoS One, 2012. **7**(6): p. e37149.

9.	Chen, Q., et al., *Rewiring of Glutamine Metabolism Is a Bioenergetic Adaptation of Human Cells with Mitochondrial DNA Mutations.* Cell Metab, 2018. **27**(5): p. 1007–1025 e5.

10.	Leung, K.Y., et al., *Partitioning of One-Carbon Units in Folate and Methionine Metabolism Is Essential for Neural Tube Closure.* Cell Rep, 2017. **21**(7): p. 1795–1808.

11.	Choi, H., et al., *Primary Cilia Negatively Regulate Melanogenesis in Melanocytes and Pigmentation in a Human Skin Model.* PLoS One, 2016. **11**(12): p. e0168025.
